# Bleb-based extravasation: conserved morphodynamics, divergent calcium control

**DOI:** 10.1101/2025.09.26.677702

**Authors:** Mizuki Morita, Manami Morimoto, Junichi Ikenouchi, Bertrand Pain, Yuji Atsuta, Yoshiki Hayashi, Takayuki Teramoto, Daisuke Saito

## Abstract

Extravasation, the exit of cells from blood vessels, is essential in both development and disease, from germ cell migration to cancer metastasis. We show that all extravasating cells examined, avian primordial germ cells and diverse human cancer lines, form Ca²⁺-dependent membrane blebs during transendothelial migration. Yet the source of Ca²⁺ diverges by lineage. PGCs and HT-1080 fibrosarcoma cells rely on store-operated Ca²⁺ entry (SOCE), whereas epithelial cancer cells such as PC-3 and MDA-MB-231 use IP₃R-mediated Ca²⁺ release from the endoplasmic reticulum. These findings establish bleb-based extravasation as a conserved morphodynamic strategy powered by distinct regulatory modules. Evolutionarily, they map onto an ancestral ER-release pathway and a metazoan-derived SOCE pathway. Conceptually, cancer blebbing emerges as a composite strategy that reactivates both ancient survival programs and developmental toolkits to maximize invasive success. This unified framework highlights Ca²⁺ supply as a critical bottleneck of vascular escape, offering new angles for targeting metastatic dissemination.

## INTRODUCTION

Extravasation, the exit of cells from blood vessels, is a fundamental process in both physiological and pathological contexts, including immune cell trafficking^1^, cancer metastasis^2^, microglial infiltration^3^, and germ cell migration^4^. In vertebrate embryos, primordial germ cells (PGCs) of avian species intravasate into the bloodstream and subsequently extravasate through the dorsal mesentery toward the gonadal region, where they initiate gametogenesis^4,5^. In metastasis, malignant cells exploit this same process to colonize distant tissues^2^. Despite its broad relevance, the morphodynamic and regulatory diversity of extravasation across lineages remains poorly understood.

Traditionally, extravasation has been attributed to actin-dependent protrusions such as lamellipodia and filopodia, which mediate endothelial transmigration through junctional openings^6^. We recently uncovered an alternative strategy: membrane blebbing. These balloon-like, pressure-driven protrusions arise from local detachment of the plasma membrane from cortical actin. Unlike lamellipodia, blebs support rapid, adhesion-independent migration and are broadly used across cell types, from amoeboid cells to mammalian immune cells^7,8^. This phenomenon was first observed *in vivo* during PGC extravasation in chick embryos^4^ and later extended to human cancer cells in the avian yolk sac vasculature (YSV)^9^. These observations provided the first *in vivo* evidence that both normal and malignant cells can deploy blebbing to breach the endothelium. Whether such protrusions are powered by a conserved molecular program or represent convergent solutions that culminate in similar morphologies remains unresolved.

Ca²⁺ signaling has been implicated in bleb initiation across multiple systems^10,11^, but whether this requirement represents a universal principle or diverges between lineages is unknown. Addressing this issue is essential, not only for defining the mechanistic logic of bleb-based migration, but also for testing whether cancer extravasation reflects co-option of developmental pathways or reactivation of more ancient cellular strategies.

Direct analysis in mammalian systems is limited by deep tissue location and rapid extravasation events. In contrast, avian embryos offer unique advantages: both the YSV and the PGC-specific extravasation vascular plexus (Ex-VaP) are planar, transparent, and accessible for single-cell *in vivo* imaging. Moreover, these models permit simultaneous transplantation of genetically distinct cells into a shared vascular environment^4,9^. This experimental platform provides an unprecedented opportunity to compare germ cells and cancer cells side by side, allowing us to dissect conserved principles and lineage-specific variations of bleb-based extravasation.

## RESULTS

### Chick PGCs form membrane blebs during extravasation

To investigate the cellular dynamics of PGC extravasation, we focused on the chick Ex-VaP at Hamburger–Hamilton (HH) stage^12^ 15, where circulating PGCs are selectively arrested prior to transendothelial migration (TEM). For live imaging, we used EGFP-expressing PGCs that had been expanded and genetically labeled *in vitro*, then reintroduced into the embryonic vasculature. Transplanted PGCs recapitulate the full migratory path to the gonads, faithfully mimicking their endogenous counterparts^4^, thus enabling high-resolution visualization under physiologically relevant conditions.

Live imaging of transplanted EGFP^+^ PGCs revealed two sequential protrusive behaviors. In the first phase, which we term *intravascular crawling*, cells rapidly produced multiple small, transient spherical protrusions (< 5 min) in random directions (Fig. 1a and Supplementary Videos1). This was followed by the emergence of a single, large, long-lived spherical protrusion (15–30 min) oriented toward the extravascular space, marking the onset of TEM (Fig. 1b and Supplementary Videos2).

**Fig. 1.**
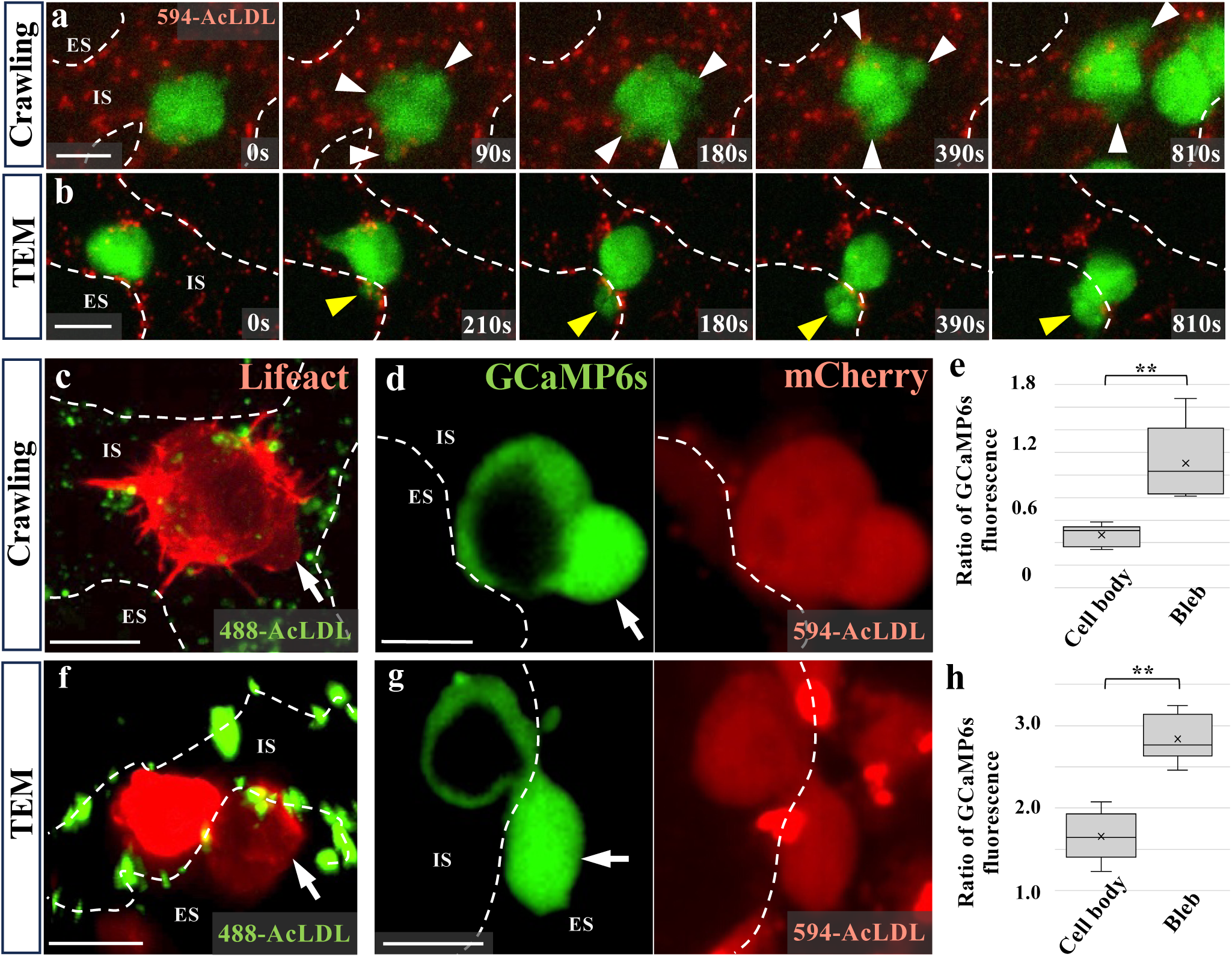
PGCs employ bleb-like protrusions during extravasation. **a, b,** Time-lapse imaging of EGFP^+^ PGCs transplanted into the Ex-VaP of HH15 chick embryos. Two distinct protrusive behaviors were observed: small transient protrusions during intravascular crawling (**a**, white arrowheads) and large, persistent protrusions during TEM (**b**, arrowheads). White dotted lines mark boundaries between intravascular space (IS) and extravascular space (ES). **c, f,** Distribution of cortical F-actin in PGC expressing Lifeact-mCherry during crawling (**c**) or TEM (**f**). Bleb-like protrusion lack cortical actin (white arrowhead). **d, g,** Ca^2+^ distribution in GCaMP6s-expressing PGC during crawling (**d**) or TEM (**g**), with co-expressed mCherry as a volume control. Protrusion shows localized Ca^2+^ enrichment (white arrowhead). **e, h,** Quantification of GCaMP6s fluorescence intensity in protrusions relative to the cell body during crawling (**e**) or TEM (**h**). N=3 embryos, respectively. Data are shown as box-and whisker plots. **P<0.01 (two-sided paired Student’s *t*-test). Scale bars, 10 μm.

To determine whether these protrusions are bona fide blebs, we examined cortical actin using Lifeact-mCherry. Both small and large protrusions showed transient loss of cortical actin (Fig. 1c,f), a hallmark of blebs^10,13–15^. We next asked whether these blebs also display localized Ca²⁺ surges, a phenomenon frequently associated with motility-driven blebbing^10,11^. Imaging with GCaMP6s revealed transient Ca²⁺ increases precisely at protrusion sites (Fig. 1d,e,g,h). Thus, PGC blebs during extravasation are defined by the combined molecular signature of actin loss and localized Ca²⁺ elevation.

Importantly, similar bleb-like protrusions were also observed in endogenous PGCs undergoing TEM *in vivo* (Extended Data Fig. 1a), confirming that blebbing is an intrinsic behavior of chick PGCs rather than an artifact of culture or transplantation.

### *In vitro* reconstitution reveals SOCE as a driver of bleb formation in chick PGCs

To dissect the mechanism of bleb formation in chick PGCs, we established a tractable *in vitro* system using under-agarose (UA) compression, a mechanical stimulus known to induce blebbing in diverse cell types^16,17^ (Extended Data Fig. 2a). Cultured PGCs reproducibly exhibited recurrent expansion and retraction of large spherical protrusions (Extended Data Fig. 2b), closely resembling their *in vivo* behavior. These protrusions coincided with localized Ca²⁺ surges and transient loss of cortical actomyosin, as visualized with GCaMP6s and Lifeact reporters (Extended Data Fig. 2c-g and Supplementary Videos3,4), consistent with bona fide membrane blebbing.

Pharmacological perturbations implicated SOCE as the source of Ca²⁺. Chelation of extracellular Ca²⁺ with BAPTA or inhibition of endoplasmic reticulum (ER) Ca²⁺ release with 2-APB^18,19^, each strongly suppressed bleb formation (Fig. 2a,b). This dual requirement pointed to store-operated Ca²⁺ entry (SOCE)^20,21^, in which ER Ca²⁺ depletion triggers Orai Ca²⁺ channel opening at the plasma membrane (PM) via ER-resident STIM sensors that translocate to ER–PM contacts^22,23^.

**Fig. 2.**
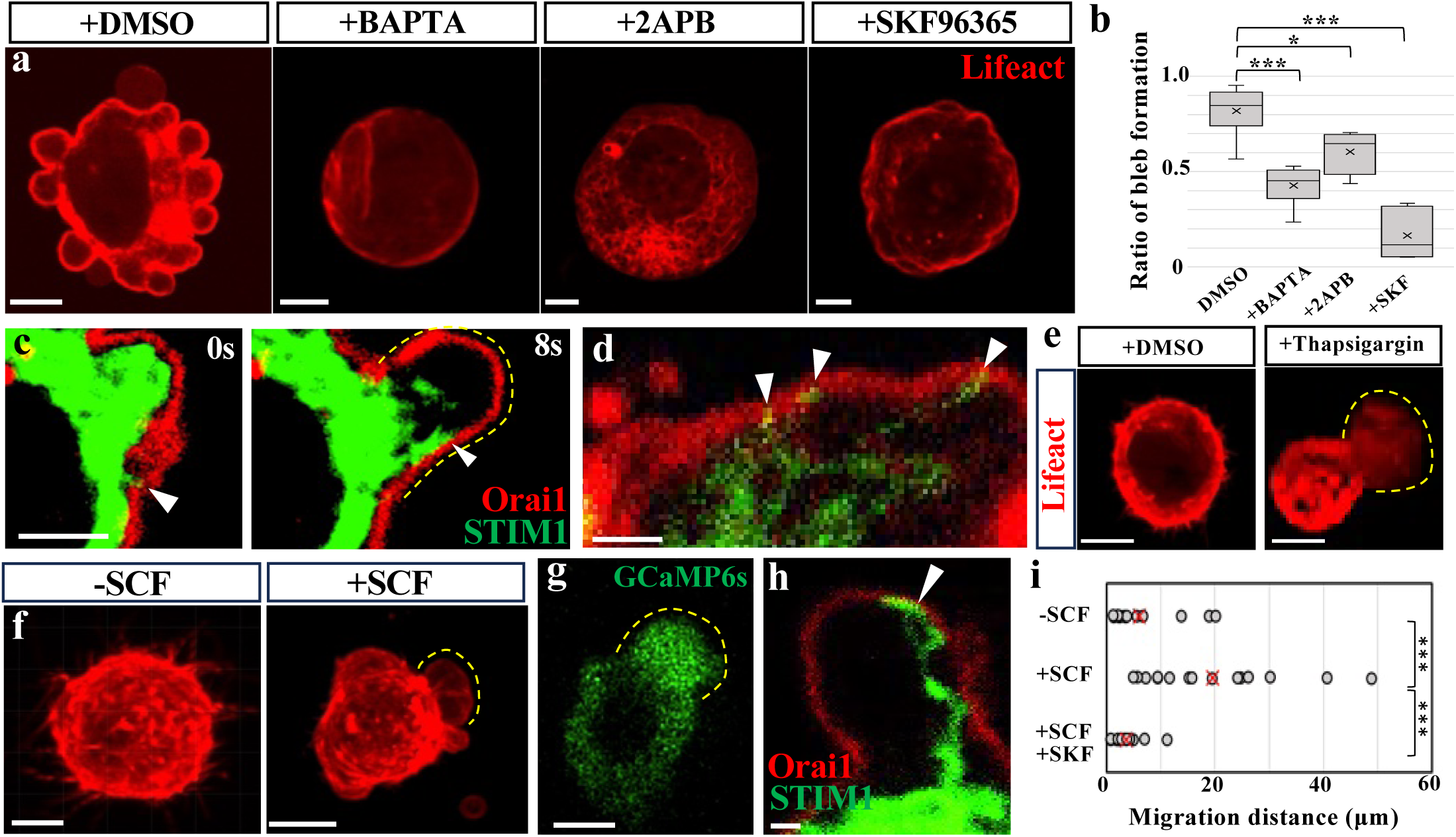
SOCE drives Ca^2+^-dependent bleb formation and migration of chick PGCs *in vitro*. **a,** Representative images of Lifeact-mCherry-expressing PGCs under agarose compression with DMSO (vehicle), extracellular Ca^2+^ chelator (BAPTA), ER Ca^2+^ release inhibitor (2-APB), or the SOCE inhibitor (SKF96365). Blebbing was strongly suppressed by BAPTA, 2-APB, and SKF. **b,** Quantification of bleb-forming area relative to total cell area (*N*=10-12 cells per group). **c, d** Time-lapse images of a PGC co-expressing STIM1-EGFP (green) and Orai1-mCheery (red), showing dynamic colocalization at ER-PM contact sites during bleb during bleb initiation and expansion (white arrowheads). **e,** PGCs cultured in Matrigel and treated with DMSO or thapsigargin, showing robust bleb induction under thapsigargin treatment. **f,** Lifeact-mCherry-expressing PGC embedded in Matrigel under SCF stimulation showing a single large bleb at the leading edge. **g**, GCaMP6s-expressing PGC showing localized Ca^2+^ accumulation at the leading bleb. **h,** Confocal images of PGCs co-expressing Stim1-EGFP (green) and Orai1-mCheery (red), demonstrating persistent colocalization at ER-PM contacts during bleb expansion (white arrowhead). **i,** Quantification of PGC migration distances over 30 minutes under the indicated conditions (*N*=15 per condition). The yellow dotted lines outline the emerging blebs. *P<0.05, ***P<0.001 (two-sided unpaired Student’s *t*-test). Scale bars, 5 μm (**a**, **e-g**); 1μm (**c, d, h**).

Re-analysis of published transcriptomic data^24^ confirmed robust expression of STIM2 and Orai1/2 in chick PGCs at HH17 (E2.5), with STIM1 also detectable at low levels (Supplementary information 1). Consistent with canonical SOCE architecture, fluorescently tagged STIM1 localized to the ER and Orai1 to the PM (Extended Data Fig. 3a,b). During bleb initiation and expansion, fluorescently tagged STIM1 and Orai1 dynamically co-localized at discrete ER–PM contacts within protrusions (Fig. 2c,d), consistent with localized SOCE activity^11^. Functionally, SOCE blockade with SKF96365 (hereafter SKF) abolished blebbing, whereas ER Ca²⁺ depletion with thapsigargin— normally sufficient to activate SOCE—robustly induced bleb formation even without mechanical stimulation (Fig. 2a,b,e). Likewise, treatment with the Ca²⁺ ionophore A23187 triggered robust blebbing in the absence of external stimuli (Extended Data Fig. 4a and Supplementary Videos5).

Together, these findings demonstrate that chick PGCs harbor the full complement of functional SOCE machinery that mediates extracellular Ca²⁺ influx to initiate and sustain bleb formation. Moreover, Ca²⁺ elevation alone is sufficient to induce blebbing *in vitro*, providing a mechanistic framework for analyzing extravasation *in vivo* and raising the question of whether the requirement for SOCE is universal or whether distinct Ca²⁺ supply routes have evolved in different lineages.

### SOCE-driven bleb formation supports PGC locomotion *in vitro*

To test whether membrane blebbing contributes to locomotion, we developed an *in vitro* assay in which chick PGCs actively migrate. We examined two candidate ligands, stromal-derived factor-1 (SDF-1) and stem cell factor (SCF), both implicated in germ cell migration^25–27^. Chick PGCs express the corresponding receptors, CXCR4 and KIT^27,28^.

When cultured alone or with COS7 cells expressing either control constructs or SDF-1, PGCs exhibited minimal motility or protrusive activity (Extended Data Fig. 5a-f). By contrast, co-culture with SCF-expressing COS7 cells robustly stimulated motility in more than half of the PGCs, although no directional migration bias was detected (Extended Data Fig. 5a-f). This system therefore provided a tractable assay to interrogate the role of SOCE in PGC migration.

Under SCF stimulation, migrating PGCs generated and maintained large balloon-like protrusions at the leading edge. Live imaging of Lifeact-mCherry and GCaMP6s-expressing cells revealed that these protrusions were actin-poor and Ca²⁺-rich (Fig. 2f,g and Extended Data Fig. 5g,h), consistent with persistent membrane blebs. For comparison, zebrafish PGCs form transient actin-brush structures during blebbing cycles^29^, whereas chick PGCs lacked such reinforcement.

Fluorescently tagged STIM1 and Orai1 co-localized at the leading edge of migrating PGCs, indicating localized SOCE activity (Fig. 2h and Extended Data Fig. 5i). Disruption of SOCE with SKF abolished bleb formation and significantly reduced migration distance (Fig. 2i and Extended Data Fig. 5j). Together, these results demonstrate that SOCE is required not only for bleb formation but also for efficient locomotion of chick PGCs *in vitro*.

### Dominant-negative Orai1 mutant cell-autonomously blocks SOCE in PGCs

To genetically dissect SOCE function *in vivo*, we established PGCs with inducible, cell-autonomous inhibition of this pathway. We used a Tet-on system^30^ to drive expression of the dominant-negative Orai1 mutant (E106Q)^11^, which abolishes Ca²⁺ conduction through Orai1 channels upon Doxycycline (Dox) administration.

In the UA compression assay, Orai1 E106Q-expressing PGCs failed to generate blebs, phenocopying the effect of pharmacological SOCE inhibition (Extended Data Fig. 6a,b). In the SCF-induced migration assay, these cells displayed severely impaired motility relative to controls (Extended Data Fig. 6c,d). Together, these results establish Orai1 E106Q as a robust genetic tool for cell-autonomous suppression of SOCE in chick PGCs, providing a basis for functional analysis *in vivo*.

### SOCE drives bleb-based extravasation and subsequent tissue migration of chick PGCs *in vivo*

To test whether SOCE-mediated bleb formation is required for PGC extravasation *in vivo*, we co-infused equal numbers of EGFP-labeled Orai1 E106Q–expressing PGCs and mCherry-labeled control PGCs into the vasculature of HH15 chick embryos (Extended Data Fig. 7a). Both populations efficiently arrested within the Ex-VaP, demonstrating that SOCE is dispensable for intravascular arrest (Extended Data Fig. 7b,c).

Live imaging 1 hour post-transplantation (hpt) revealed a marked defect in subsequent behaviors: Orai1 E106Q^+^ PGCs failed to initiate blebbing, remained spherical, and displayed neither intravascular crawling nor TEM (Fig. 3a–c and Extended Data Fig. 7d and Supplementary Videos6). To quantify extravasation efficiency, we transplanted the same cell combinations into HH15 quail embryos and stained for the endothelial marker QH1 at 6 hpt. Quail were used for this analysis because QH1 provides clear and specific endothelial labeling^5,31^, while comparable antibodies are not available for chick, ensuring acute discrimination between intra- and extravascular PGCs^9^. Orai1 E106Q^+^ PGCs extravasated at ∼40% lower efficiency than co-infused control PGCs (Fig. 3d,e). By contrast, co-transplanted EGFP^+^ and mCherry^+^ controls showed comparable efficiencies, excluding labeling or tracking artifacts (Fig. 3d,e).

**Fig. 3.**
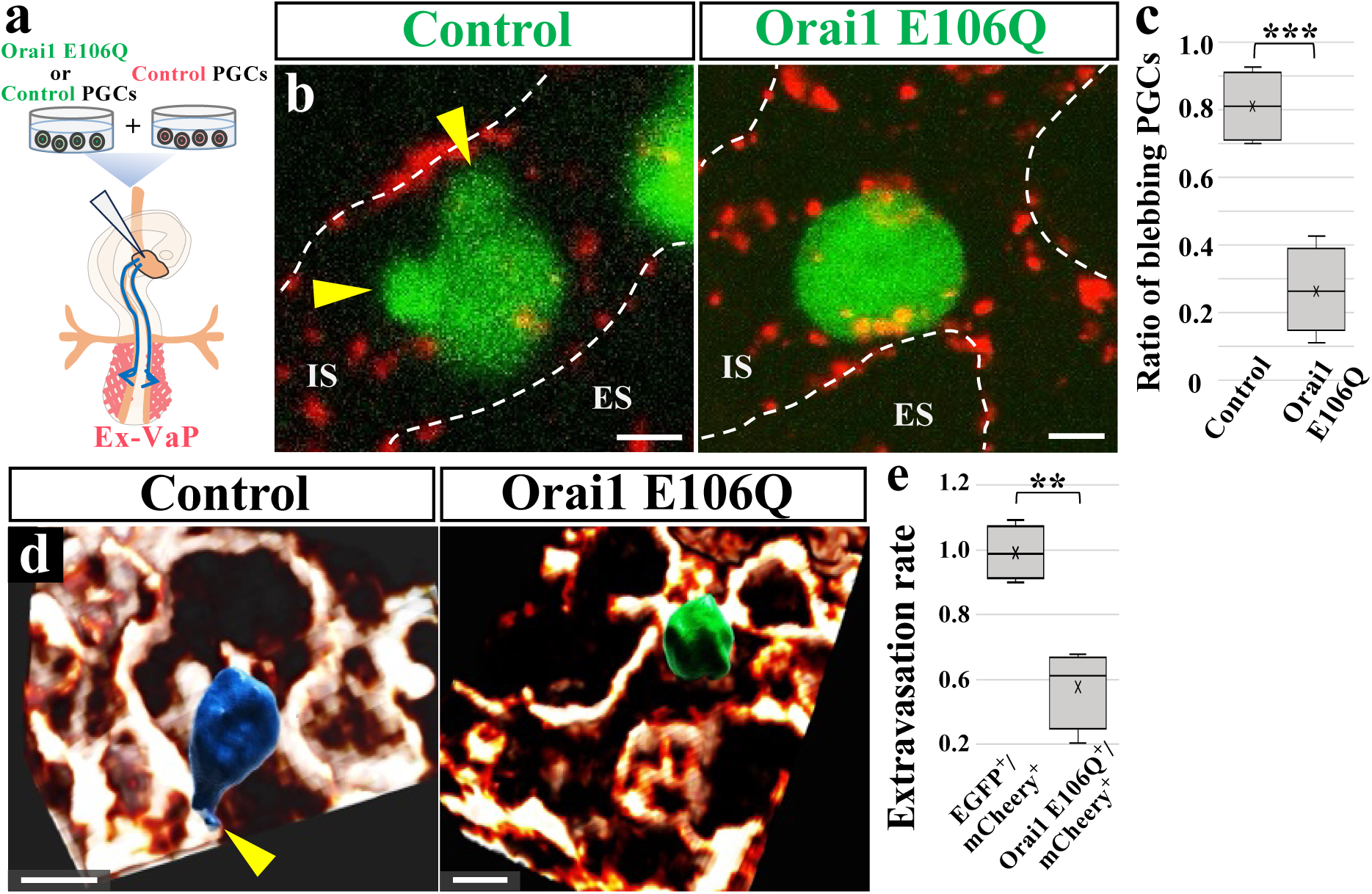
SOCE is required for PGC extravasation. **a,** Schematic of the co-infusion strategy introducing control or Orai1 E106Q-expressing PGCs, together with an equal number of mCherry^+^ PGCs as internal controls, into the vasculature of HH15 chick or quail embryos. **b**, Representative images of control and Orai1 E106Q^+^ PGCs arrested in the Ex-VaP vasculature. Endothelium is labeled with 594-AcLDL (red). Yellow arrowheads indicate bleb formation in a control cell. White dashed lines outline intravascular (IS) and extravascular (ES) compartments. **c**, Quantification of protrusion-forming PGCs (*N*=4 embryos per group). **d,** 3D reconstructions of the Ex-VaP region in HH16 quail embryos at 6 hpt. Vascular network is labeled by QH1 (brown). A PGC is labeled with mCherry (blue) and EGFP (green). Left: ventral view; right: transverse view. The yellow arrowhead indicates the leading bleb of the transmigrating PGC. **e,** Quantification of extravasation efficiency for Orai1 E106Q^+^ and control PGCs, normalized to co-infused mCherry^+^ controls (*N*=4 embryos). **P<0.01, ***P<0.001 (two-sided unpaired Student’s *t*-test). Scale bars: 5 μm (**b, d left**), 10 μm (**d right**).

We next examined whether SOCE is also required for post-extravasation migration. At E4.5 (2 days post-transplantation), many control PGCs had reached the dorsal mesentery or gonadal primordium, whereas significantly fewer Orai1 E106Q+ PGCs were present in these regions (Extended Data Fig. 7e,f). The relative abundance of mutant cells decreased from 63% in the mesentery to 22% in the gonad, consistent with a progressive defect in tissue migration. Importantly, *in vitro* assays confirmed that SOCE inhibition did not affect PGC survival or proliferation (Extended Data Fig. 7g,h). Together, these results establish that SOCE is essential for bleb initiation, vascular exit, and subsequent migration toward the gonad, underscoring its central role in the *in vivo* trajectory of chick PGCs.

### SOCE drives bleb-based transendothelial migration of HT-1080 human cancer cells

Given that SOCE governs membrane blebbing and extravasation in chick PGCs, we asked whether the same mechanism also operates in human cancer cells. Aggressive cancer cells such as HT-1080 fibrosarcoma cells migrate and invade to form metastatic tumors, making them a relevant model for testing whether bleb-based extravasation contributes to malignant dissemination^32^. When transplanted into the vasculature of HH15 avian embryos, HT-1080 cells rapidly arrested in the YSV within 20 minutes, without evidence of intravascular crawling, and initiated TEM by extending thin pseudopodia, forming large bleb-like protrusions, or combining both modes into mixed-type leading edges (Fig. 4a and Extended Data Fig. 8a,b). These observations are consistent with previous reports indicating that ∼10% of extravasating cancer cells employ bleb-driven mechanisms^9^.

**Fig. 4.**
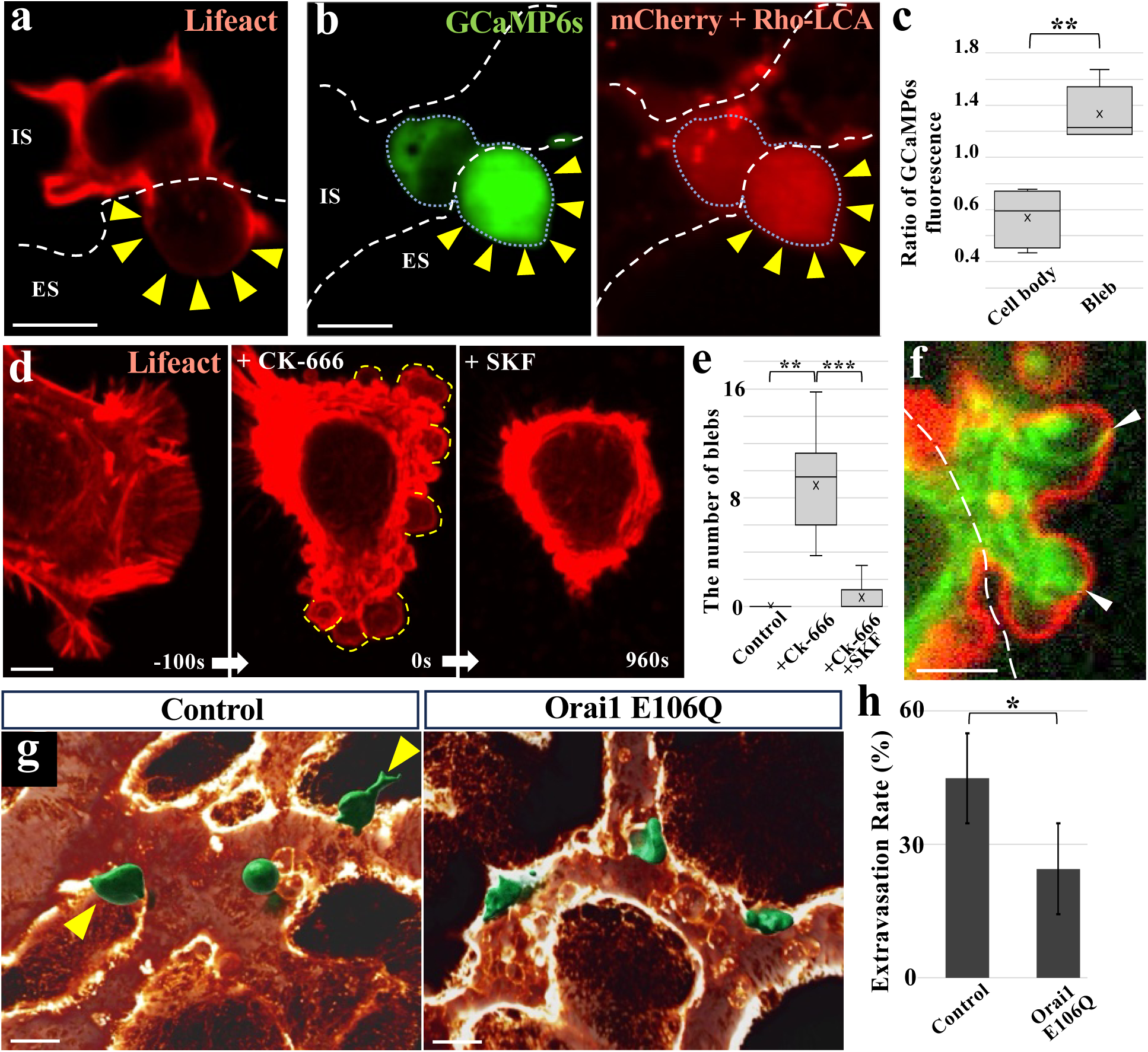
SOCE drives bleb-based TEM of HT-1080 human cancer cells. **a,** Representative image of an HT-1080 cell undergoing TEM in the YSV. Lifeact-mCherry reveals actin-poor blebs (yellow arrowheads). **b,** Left: GCaMP6s-expressing HT-1080 cell showing localized Ca^2+^ accumulation within a protrusion (yellow arrowheads). Right: merged image with mCherry and Rho-LCA. **c,** Quantification of GCaMP6s fluorescence intensity in blebs relative to the cell body in transmigrating HT-1080 cells (*N*=3 embryos). **d,** Time-lapse imaging of an HT-1080 cell exposed to CK-666 (Arp2/3 inhibitor) to induce blebbing (yellow dotted lines), followed by SKF96365 in 2D culture. CK-666-induced blebs are suppressed after SOCE inhibition. **e,** Quantification of the proportion of bleb-forming cells before and after SKF96365 treatment (*N*=8 cells). **f,** Confocal image of an HT-1080 cell co-expressing STIM1-EGFP (green) and mCherry-CAAX (red) during TEM. White arrowheads indicate contact sites of STIM1 and the PM within a bleb. **g,** 3D reconstructions of control (left) and Orai1 E106Q-expressing (right) HT-1080 cells (green) undergoing TEM in the YSV at 3.5 hpt. Vasculature is visualized with Rho-LCA (brown). Control cells exhibit active TEM (yellow arrowheads), whereas Orai1 E106Q cells frequently remain intravascular. **h,** Quantification of extravasation efficiency for control and Orai1 E106Q-expressing HT-1080 cells in the YSV at 3.5 hpt (N=4 embryos; 102 and 130 cells analyzed, respectively). White dashed lines outline IS and ES. **c and e:** **P<0.01, ***P<0.001 (two-sided, paired Student’s *t-*test), **h**: *P<0.05 (two-sided, unpaired Student’s *t-*test). Scale bars: 10 μm (**a, b**); 5 μm (**d**, **f**)**; 30** μm (**g**). Error bars indicate s.e.m.

To determine whether these protrusions are bona fide blebs, we imaged HT-1080 cells expressing GCaMP6s and Lifeact-mCherry during TEM. Balloon-like protrusions exhibited localized cytoplasmic Ca²⁺ surges and depletion of cortical F-actin (Fig. 4a–c), recapitulating the molecular hallmarks of blebs observed in chick PGCs.

We next assessed the requirement for SOCE. Inhibition of Arp2/3-mediated actin polymerization with CK-666 induces spherical bleb-like protrusions^33^, which were abolished by the SOCE inhibitor SKF (Fig. 4d,e and Supplementary Videos7), indicating that SOCE activity is necessary for bleb formation under these conditions. Consistently, thapsigargin treatment also induced robust blebbing in 2D culture (Extended Data Fig. 8c,d), providing additional support for SOCE involvement.

During TEM, GFP-STIM1 accumulated at the plasma membrane within bleb-like protrusions (Fig. 4f), consistent with ER–PM contact formation and localized SOCE activation. To functionally test the role of SOCE *in vivo*, we generated HT-1080 cells stably expressing Orai1 E106Q. Upon vascular transplantation, Orai1 E106Q-expressing cells displayed normal arrest and protrusion formation (Extended Data Fig. 8e-h), but showed a significantly reduced rate of TEM compared to controls (Fig. 4g,h and Extended Data Fig. 8i). Finally, treatment with the Ca²⁺ ionophore A23187 was sufficient to induce robust blebbing in HT-1080 cells in 2D culture (Extended Data Fig. 4b and Supplementary Videos8). Together, these results establish that SOCE-dependent Ca²⁺ entry from the extracellular space is indispensable for bleb-driven extravasation in HT-1080 cells. This parallels the mechanism observed in chick PGCs and highlights SOCE as a conserved driver of bleb-based TEM across both development and cancer.

### IP₃R-mediated Ca²⁺ release drives SOCE-independent bleb formation and extravasation in PC-3 and MDA-MB-231 cancer cells

While SOCE-dependent bleb formation was essential for TEM in both chick PGCs and HT-1080 cells, it remained unclear whether this mechanism was conserved in other metastatic cancer types. To address this, we analyzed PC-3 (prostate cancer) and MDA-MB-231 (breast cancer) cells, which also form bleb-like protrusions during extravasation in the avian model^7^. Similar protrusions have been reported under culture conditions^34,35^.

Consistent with bona fide blebs, these balloon-like protrusions were depleted of cortical actin (Fig. 5a). Because GCaMP6s expression impaired viability in these epithelial cancer cells, we instead used chemical indicators to probe Ca²⁺ dynamics. In 2D culture, spontaneously blebbing cells exhibited higher Ca²⁺ signals within blebs compared to the adjacent cell body, as revealed by Fluo-4 AM imaging (Fig. 5b,c).

**Fig. 5.**
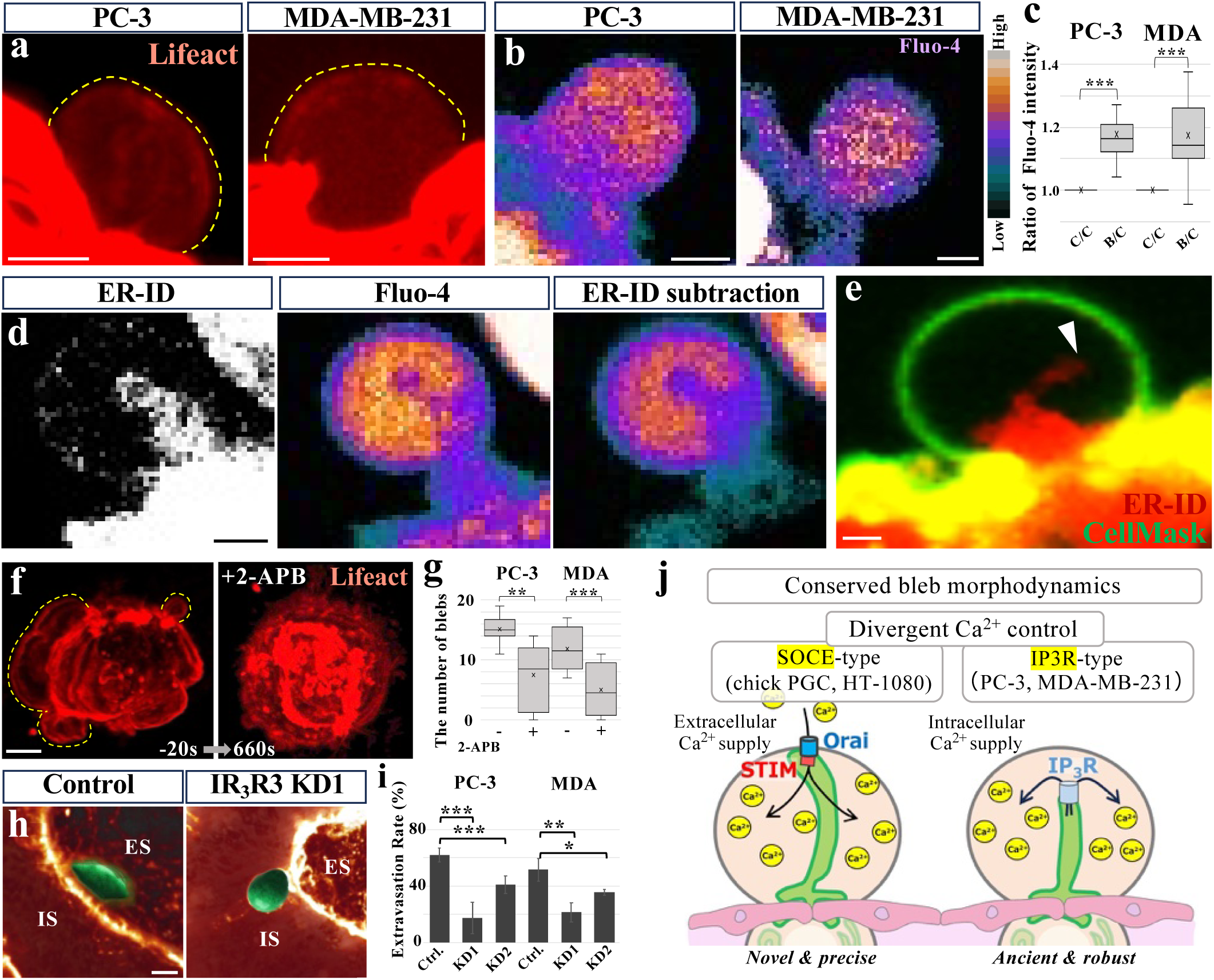
IP₃R-mediated Ca^2+^ release drives SOCE-independent bleb formation and TEM in epithelial cancer cells. **a,** Lifeact-mCherry-expressing PC-3 and MDA-MB-231 cells during bleb formation (yellow dotted lines). **b,** Fluo-4-loaded PC-3 and MDA-MB-231 cells during bleb formation. **c,** Quantification of Fluo-4 fluorescence intensity in blebs relative to the cell body (*N*=8 cells per group). Ratios are shown as C/C (cell body-to-cell body, control) and B/C (bleb-to-cell body). **d,** Dual imaging of ER-ID and Fluo-4 in an MDA-MB-231 cell during bleb formation. Left and middle: raw images; right: subtraction of ER–ID signal from Fluo-4. **e,** Dual visualization of CellMask (green) and ER-ID (red) showing ER tubules extending into a bleb of an MDA-MB-231 cell (white arrowhead). **f,** Time-lapse imaging of a PC-3 cell before and after 2-APB addition in 2D culture. Time is shown in seconds relative to the initial addition of 2-APB. The yellow dotted lines outline the emerging blebs. **g,** Quantification of the number of blebs before and after 2-APB treatment in PC-3 and MDA-MB-231 (*N*=8 and 6 cells, respectively). **h,** 3D reconstructions of control and IP_3_R3 KD1 MDA-MB-231 cell (green iso-surfaces) in the YSV at 3.5 hpt. Blood vessels were labeled by intravenous injection of Rho-LCA (brown). Endothelial volumes were digitally clipped to visualize transmigrating cells. **i,** Quantification of extravasation efficiency for control, IP_3_R3 KD1, and IP_3_R3 KD2 PC-3 cells and MDA-MB-231 cells. Efficiency was calculated as the percentage of cells located outside the vasculature among all labeled cells in the YSV (*N*=5 and 4 embryos, respectively). **j,** Schematic of SOCE-type and IP_3_R-type pathways. Bleb-based extravasation is conserved in form but divergent in Ca^2+^ supply. *P<0.05, **P<0.01, ***P<0.001. (**c, i**: Two-sided Welch’s t-test; **g**: two-sided paired Student’s *t-*test). Scale bars: 2μm in **a-f**, 10 μm in **h**. Error bars indicate s.e.m.

To test whether Ca²⁺ elevation per se could drive blebbing, we applied the ionophore A23187. As in PGCs and HT-1080 cells, A23187 was sufficient to robustly induce bleb formation in both PC-3 and MDA-MB-231 cells (Extended Data Fig. 4c,d and Supplementary Videos9,10), confirming that Ca²⁺ is a conserved driver of bleb initiation. However, manipulation that abolished SOCE-dependent blebbing in PGCs and HT-1080 cells failed to affect PC-3 and MDA-MB-231 cells. Pharmacological SOCE inhibition with SKF had no effect on bleb formation (Extended Data Fig. 9a–c and Supplementary Videos11,12). Depletion of ER Ca²⁺ stores with thapsigargin, which normally activates SOCE, likewise failed to induce blebbing (Extended Data Fig. 9d,e). Expression of dominant-negative Orai1 E106Q also left TEM efficiency unchanged in both PC-3 and MDA-MB-231 cells, in stark contrast to the impairment seen in HT-1080 cells (Extended Data Fig. 9f–h). These results demonstrate that PC-3 and MDA-MB-231 cells do not rely on SOCE. Notably, the persistence of blebbing under extracellular Ca²⁺ chelation with BAPTA suggests the involvement of an alternative, intracellular Ca²⁺source (Extended Data Fig. 9i–k and Supplementary Videos13,14).

To directly test this possibility, we examined the spatial relationship between ER and Ca²⁺ dynamics. Strikingly, dual visualization with Fluo-4 and ER-ID revealed that ER tubules extend into blebs, and localized Ca²⁺ elevations consistently arose in close proximity to these ER structures (Fig. 5d and Extended Data Fig. 10a). Importantly, these ER extensions did not appear to reach or stabilize at the PM, in contrast to the ER-PM junctions characteristic of SOCE-dependent blebs (Fig. 5e and Extended Data Fig. 10b). This indicates that, rather than engaging in SOCE, these epithelial cancer cells mobilize ER-derived Ca²⁺ directly within blebs to fuel their expansion.

Consistent with this model, pharmacological inhibition of IP₃Rs, a major class of ER Ca²⁺ release channels, with 2-APB markedly reduced blebbing (Fig. 5f,g and Extended Data Fig. 10c and Supplementary Videos15,16). To further test the role of IP₃R signaling, we knocked down IP₃R3, the predominant isoform in both cell lines^36^. IP₃R3 knockdown significantly reduced the frequency of blebs and other protrusions *in vitro* (Extended Data Fig. 10d). *In vivo*, IP₃R3-deficient cells exhibited normal vascular arrest but markedly reduced extravasation efficiency (Fig. 5h,i and Extended Data Fig. 10h–m), implicating IP₃R3 in promoting TEM through its control of Ca²⁺-dependent protrusive activity.

Together, these findings define a distinct, SOCE-independent mechanism in which IP₃R3-mediated Ca²⁺ release from ER reservoirs drives extravasation in PC-3 and MDA-MB-231 cells. This establishes a lineage-specific divergence of Ca²⁺ supply routes: SOCE operates in chick PGCs and HT-1080 cells, whereas IP₃R3-mediated release predominates in PC-3 and MDA-MB-231 cells (Fig. 5j).

## DISCUSSION

Our study establishes a unified framework to directly compare PGCs and cancer cells within the same vascular context of *in vivo* extravasation. Using this system, we show that, despite differences in frequency, both lineages employ membrane blebbing to breach the endothelium. Blebs are particularly suited for TEM because, unlike lamellipodia or filopodia that rely on substrate adhesion and actin polymerization, they are pressure-driven protrusions that transiently detach the actin cortex from the plasma membrane ^7^. This mechanism enables cells to forcibly push through narrow endothelial gaps with strong propulsive forces even under confinement. One of our central findings is that Ca²⁺ elevation is universally sufficient to trigger bleb formation, highlighting Ca²⁺ as a conserved driver of extravasation morphodynamics. Yet, Ca²⁺ supply diverges: PGCs and HT-1080 fibrosarcoma cells rely on SOCE-mediated influx from the extracellular milieu, whereas PC-3 and MDA-MB-231 epithelial cancer cells depend on IP₃R3-mediated Ca²⁺ release from the ER. This divergence delineates two mechanistic classes of bleb-based migration: an extracellular Ca²⁺-driven SOCE module and an intracellular ER-release module.

The existence of two distinct Ca²⁺ supply modules suggests that blebs can be tuned for different functional demands. SOCE provides precision, suited for lineages requiring localized, directional blebs such as PGCs navigating to the gonad. A parallel can be drawn from immune cells, where SOCE supplies spatially restricted Ca²⁺ signals at the immunological synapse to ensure precise activation^37^. By contrast, IP₃R-mediated release provides robustness by ensuring bleb initiation even under fluctuating extracellular Ca²⁺ conditions. In PC-3 and MDA-MB-231 cells, this is achieved through localized Ca²⁺ elevations at ER extensions that penetrate into blebs, a mode of regulation distinct from the ER-PM junction-based SOCE. Such ER-driven supply confers robustness, enabling epithelial cancers to prioritize persistence over precision while maintaining reliable protrusion under variable environments (Fig. 5j)^38,39^.

From an evolutionary perspective, ER-based release represents the ancestral mode: Dictyostelium lacks STIM/Orai^40^ yet blebs through an IP₃R-like channel (IplA)^41–43^. SOCE represents a later innovation, restricted to metazoans^43,44^, that enables spatially precise Ca²⁺ delivery through ER–PM nanodomains. The coexistence of these two modules reveals that cancer does not simply revert to a single primitive state but instead reactivates multiple layers of cellular evolution. IP₃R-based release embodies atavism, the reawakening of ancient unicellular programs optimized for robustness under fluctuating conditions. In contrast, SOCE-dependent control exemplifies developmental co-option, where cancer appropriates the finely tuned morphodynamics of embryonic migration to achieve precision and directionality. Thus, cancer blebbing should be viewed as a composite strategy, simultaneously drawing on ancestral survival mechanisms and developmental toolkits to maximize invasive success (Fig. 5j)^45–48^.

Zebrafish PGCs exemplify an intermediate state between these two modules. In this system, blebs are triggered by localized intracellular Ca²⁺ elevations^10^, but the precise source of Ca²⁺ remains unresolved. Previous work has shown that expression of constitutively active STIM1 can induce ectopic Ca²⁺ elevations and bleb formation^10^, demonstrating that SOCE can be harnessed in these cells. At the same time, the physiological requirement for SOCE under normal developmental conditions has not been established, leaving open the possibility that IP₃R-dependent release also contributes. Thus, zebrafish PGCs appear to retain both ancestral IP₃R-type and derived SOCE-type potentials, positioning them as a “missing link” that bridges the evolutionary diversification of Ca²⁺ supply modules. More broadly, PGCs across species reveal a continuum from ancestral to derived Ca²⁺ strategies, underscoring how developmental germ cells can illuminate the evolutionary layering of bleb regulation.

Clinically, this bifurcation suggests that lineage-specific Ca²⁺ supply routes could be directly leveraged for intervention. SOCE inhibitors, several of which are already under preclinical evaluation^49^, may selectively target lineages that rely on SOCE-dependent blebbing (such as chick PGCs and fibrosarcoma cells), while IP₃R blockade could constrain bleb-based extravasation in epithelial cancer models. Our conclusions regarding epithelial cancers are based on PC-3 and MDA-MB-231 cells, and it remains to be determined whether this IP₃R-dependent mode is a broader hallmark of epithelial malignancies, particularly adenocarcinomas, or a lineage-specific adaptation. Because these modules are mechanistically distinct, their activity may also serve as a biomarker to stratify tumors according to their invasive strategies and to guide therapeutic choices. More broadly, defining the Ca²⁺ supply logic of extravasation opens the possibility of designing interventions that block metastatic dissemination at its vascular bottleneck, a stage that remains largely untargeted in current oncology.

## MATERIALS and METHODS

### Animals, staging, and animal care

Fertilized chicken (*Gallus gallus domesticus*, White leghorn) eggs and fertilized quail (*Coturnix japonica*) eggs were purchased from Yamagishi poultry farm (Mie, Japan) and from Nagoya University through the National Bio-Resource Project of the MEXT, Japan, respectively. Eggs were incubated at 38.5℃, and embryos were staged by Hamburger and Hamilton’s stage^12^. All animal experiments were performed with the approval of the Institutional Animal Care and Use Committees at Kyushu University.

### Plasmid constructions

pT2A-BI-Gap-TdTomato-TRE: pT2AL200R150G vector^50^ was digested with XhoI-BglII. These sites were blunt-ended, and inserted with the fragment of pBI-TRE (Clontech) containing the bidirectional tetracycline-responsive element (TRE) with two minimal promoters of CMV in both directions, and two polyA-additional sequences of the rabbit beta globin gene. This vector was designated as pT2A-BI-TRE. The full-length of Gap-TdTomato^51^ was amplified by PCR, and subcloned into the SpeI-HindIII site of pT2A-BI-TRE. pT2A-BI-Gap-TdTomato-TRE-(SCF or SDF-1): The ORF of chicken SCF or chicken SDF-1^52^ was subcloned into the MluI-EcoRV site of pT2A-BI-Gap-TdTomato-TRE. The ORF of SCF was amplified and isolated from cDNA derived from E2.5 whole chick embryo by PCR. pT2A-BItight-EGFP-TRE: pT2AL200R150G vector was digested with XhoI-BglII. These sites were blunt-ended, and inserted with the fragment of pBItight-TRE (Clontech) containing the improved bidirectional tetracycline-responsive element (TRE) with two minimal promoters of CMV in both directions, and two polyA-additional sequences of the rabbit beta globin gene. This vector was designated as pT2A-BItight-TRE. The full-length of EGFP was amplified by PCR, and subcloned into the EcoRI-PstI site of pT2A-BItight-TRE. pT2A-BItight-EGFP-TRE-Orai1 E106Q: The ORF of human Orai1 with E106Q mutation and myc tag (addgene #22754) was amplified by PCR, and subcloned into the MluI-EcoRV site of pT2A-BItight-EGFP-TRE. pT2A-BItight-mCherry-TRE: The full-length of mCherry was amplified by PCR, and subcloned into the EcoRI-BglII site of pT2A-BItight-TRE. pT2A-BItight-GCaMP6s-2AP-mCherry-TRE: The consecutive sequences of GCaMP6s-2AP-mCherry^53^ was amplified by PCR, and subcloned into the EcoRI site of pT2A-BItight-TRE by In-Fusion^®^ HD cloning kit (TaKaRa). pT2A-BItight-Lifeact-mCherry-TRE: Lifeact-mCherry was amplified by PCR using pT2A-BI-TRE-LAP2bcGFP-Lifeact-mCherry^9^ as a template, and subcloned into the MluI-EcoRV site of pT2A-BItight-TRE. pT2A-CAGGS-EGFP-CAAX-IRES2-PuroR: The sequences of IRES2 and PuroR were amplified by PCR, and subcloned into the XhoI-EcoRI site and EcoRI-BglII site into pT2A-CAGGS, respectively. This vector was designated as pT2A-CAGGS-IRES2-PuroR. The full-length of EGFP-CAAX was amplified by PCR, and subcloned into the PstI-XhoI site of pT2A-CAGGS-IRES2-PuroR. pT2A-CAGGS-MRLC1-EGFP-IRES-PuroR: MRLC1-EGFP was amplified by PCR, and subcloned into the NotI-XhoI site of pT2A-CAGGS-IRES2-PuroR. pT2A-CAGGS-Tet3G-IRES2-PuroR: The full-length of Tet3G was amplified by PCR, and subcloned into the NotI-XhoI site of pT2A-CAGGS-IRES2-PuroR. pT2A-BItight-mCherry-CAAX-TRE: mCherry-CAAX was amplified by PCR, and subcloned into the BglⅡ site and EcoRV site of pT2A-BItight-TRE. pT2A-BItight-stim1-EGFP-TRE: Stim1-EGFP was amplified by PCR (addgene #210370), and subcloned into the MluI-EcoRV site of pT2A-BItight-TRE.

Primer sequences are as described below.

Gap43-SpeI-F: agaactagtggatccccgcggATGTTGTGCTGTATGAGAAGAACCAAGC

TdTomato-HindIII-R: gacaagcttgaattcTTACTTGTACAGCTCGTCCATGCCGTACAG

SDF-1-MluI-F: cagacgcgtATGGACCTCCGCGCCCTGGCTCTGCTCGCCT

SDF-1-EcoRV-R: agagatatcTTACTTGTTTAAAGCTTTCTCCAGATATTCC

SCF-MluI-F: aattacgcgtATGAAGAAGGCACAAACTTGGATTATCAC

SCF-EcoRV-R: aattgatatCTACACTTGTAGATGTTCTTTTTCTTTTTGCTGCAACA

EGFP-EcoRI-F: ggagaattcATGGTGAGCAAGGGCGAGGAGCTGTTCACCG

EGFP-PstI-R: agactgcagTTACTTGTACAGCTCGTCCATGCCGAGAGT

Orai1 E106Q-MluI-F: ctgacgcgtACCATGCATCCGGAGCCCGCCCCGCCCC

Orai1 E106Q-EcoRV-R: ggagatatcCTAGAGATCTTCCTCAGAAATGAGC

mCherry-EcoRI-F: ggagaattcATGGTGAGCAAGGGCGAGGAGGATAACATGG

mCherry-BglII-R: caggagatctTTACTTGTACAGCTCGTCCATGCCGCCGGT

GCaMP6s-inf-F: gtcagatcgcctggaATGGGTTCTCATCATCATCATCATCATGGTATGG

mCherry-inf-R: gactgcagcctcaggagatcTTACTTGTACAGCTCGTCCATGCCGCCGG

Gap43-MluI-F: attacgcgtcggcaccATGCTGTGCTGTATGAGAAGAACCAAGCAGGTG

mOrange-EcoRV-R: tctgataTCACTTGTACAGCTCGTCCATGCCGCCG

stim1-EGFP-MluI: cgatacgcgtATGGATGTATGCGTCCGTCT

stim1-EGFP-EcoRV: tggagatatcTTACTTGTACAGCTCGTCCA

IRES2-XhoI-F: aatctcgagGCCCCTCTCCCTCCCCCCCCCCTAAC

IRES2-EcoRI-R: aatgaattcTGTGGCCATATTATCATCGTGTTTTTCA

PuroR-EcoRI-F: agagaattcATGACCGAGTACAAGCCCACGGTGCGCCTCG

PuroR-BglII-R: aaaagatctTCAGGCACCGGGCTTGCGGGTCATGCACCAGGT

EGFP-PstI-F: aagctgcagcggcaccATGGTGAGCAAGGGCGAGGAGCTGTTCACCGGG

CAAX-XhoI-R: gcctcgagTTACATAATTACACACTTTGTCTTTGACTTCTTTTTCTTCT

MRLC1-NotI-F: tgcagcggccgcATGTCCAGCAAGCGGGCCAA

EGFP-XhoI-R: gggcctcgagTTACTTGTACAGCTCGTCCATG

Tet3G-NotI-F: gcagcggccgcATGTCTAGACTGGACAAGAGCAAAGTC

Tet3G-XhoI-R: ggcctcgagTTACCCGGGGAGCATGTCAAGGTCAAAAT

mCherry-EcoRI-F: ggagaattctagatatcggcaccATGGTGAGCAAGGGCGAGGAGGATAA

CAAX-BglII-R: aggagatcTTTACATAATTACACACTTTGTCTTTGACTTCTTTTTCTTC

### Establishment and maintenance of PGCs in culture

Circulating PGCs along with blood cells were harvested from blood of HH 15 chicken embryos, and were cultured in Ca²⁺-free DMEM (Gibco) diluted with water, containing <u>F</u>GF2 (Wako), <u>A</u>ctivin A (APRO Science), and <u>c</u>hicken <u>s</u>erum (FAcs medium) according to the method previously described ^54,55^. After one month, expanded PGCs were cryo-preserved at -80℃ in Bambanker (NIPPON Genetics) until used for experiments.

### Maintenance of HT-1080, PC-3, and MDA-MB-231 in culture

HT-1080 cells (JCRB Cell Bank) and MDA-MB-231 cells (JCRB Cell Bank) were maintained in Dulbecco’s modified Eagle’s medium (DMEM; Nakarai tesque) containing 4.5g/l Glucose, L-Glutamine, and sodium pyruvate, supplemented with 10% (v/v) fetal bovine serum (FBS; Gibco), GlutaMax^TM^ (Gibco), and 1% (v/v) penicillin-streptomycin-amphotericin B (PSA; FUJIFILM Wako Pure Chemical Corporation).

PC-3 cells (ATCC) were cultured in Roswell Park Memorial Institute medium 1640 (RPMI1640; Wako) supplemented with 10% (v/v) FBS (Gibco), GlutaMax^TM^ (Gibco), and 1% (v/v) PSA.

All cells were incubated at 37°C in a humidified atmosphere of 5% CO_2_ and passaged using 0.05% trypsin-EDTA (Gibco).

### Plasmid transfection and establishment of gene-manipulated PGC lines and cultured cell lines (COS7, HT-1080, PC-3, MDA-MB-231)

#### PGCs

Cultured PGCs (5 x 10^4^) were washed with OPTI-MEM (Gibco) and seeded into 96-well plates containing 100 μl of FAot medium^55^ (supplemented with ovo-transferrin instead of chicken serum) without heparin and PSA for 3 h. A transfection mixture containing 0.2 μg of total plasmid DNA and 0.5 μl of Lipofectamine 2000 (ThermoFisher Sciencetific) in 50 μl of OPTI-MEM was then added. After 6 h, the medium was replaced with conventional FAcs medium containing heparin and antibiotics.

Seven plasmid sets were used for PGC transfection:

pT2A-BItight-GCaMP6s-2AP-mCherry + pT2A-CAGGS-Tet3G-IRES2-PuroR + pCAGGS-T2TP^56^

pT2A-BItight-Lifeact-mCherry-TRE + pT2A-CAGGS-Tet3G-2AP-PuroR^4^ + pCAGGS-T2TP

pT2A-BItight-EGFP-TRE + pT2A-CAGGS-Tet3G-IRES2-PuroR + pCAGGS-T2TP

pT2A-BItight-mCherry-TRE + pT2A-CAGGS-Tet3G-IRES2-PuroR + pCAGGS-T2TP

pT2A-BItight-EGFP-TRE-Orai1 E106Q + pT2A-CAGGS-Tet3G-IRES2-PuroR + pCAGGS-T2TP

pT2A-BItight-stim1-EGFP-TRE + pT2A-BItight-mCherry-CAAX-TRE + pT2A-CAGGS-Tet3G-IRES2-PuroR + pCAGGS-T2TP

pT2A-CAGGS-MRLC1-EGFP-IRES-PuroR + pCAGGS-T2TP

Following transfection, PGCs were cultured in FAcs medium supplemented with 0.1-0.3 μg/ml puromycin for 3 days. In addition, plasmids encoding Orai1-mCherry (pCAGGS-neo vector) and EGFP-tagged STIM1 (pCAGGS-neo vector)^11^ were introduced using the same method, and cells were used for experiments one day later.

#### COS7 cells

COS7 cells were maintained in Dulbecco’s modified Eagle’s medium (DMEM) supplemented 1.5g/L sodium bicarbonate, 10% fetal bovine serum (FBS), 50 IU/ml penicillin and 50mg/ml streptomycin at 37℃. Cells were transfected with following sets using Lipofectamine 2000 according to the manufacture’s instruction:

pT2A-BI-GapTdTomato-TRE + pT2K-M2-IRES2-NeoR^52^ + pCAGGS-T2TP

pT2A-BI-GapTdTomato-TRE-SCF + pT2K-M2-IRES2-NeoR + pCAGGS-T2TP

pT2A-BI-GapTdTomato-TRE-SDF-1 + pT2K-M2-IRES2-NeoR + pCAGGS-T2TP

Stable lines were selected in G418-containing medium.

#### HT-1080 cells (lipofection)

HT-1080 cells (1 × 10^6^) were washed with OPTI-MEM and seeded into 6-well plates (1 ml of medium without PSA) for 18-24 h (overnight). A transfection mixture (4.0 μg of total plasmid DNA + 7.0 μl of Lipofectamine 2000 in 250 μl of OPTI-MEM) was added. Two plasmid sets were used:

pT2A-BItight-Lifeact-mCherry-TRE + pT2A-CAGGS-Tet3G-2AP-PuroR + pCAGGS-T2TP

pT2A-BItight-EGFP-TRE + pT2A-CAGGS-Tet3G-2A-PuroR + pCAGGS-T2TP

12 hours later, the medium was replaced with conventional medium containing antibiotics. After 2 days, cells were cultured in 2-5 μg/ml puromycin for 2 days. EGFP-positive cells were then sorted using an SH800 cell sorter (Sony).

#### HT-1080 cells (electroporation)

HT-1080 cells (1 × 10^6^) were also electroporated using the Neon Transfection System (Invitrogen). For electroporation, 6 × 10^4^ cells were transfected with:

pT2A-BItight-stim1-EGFP-TRE + pT2A-BItight-mCherry-CAAX-TRE + pT2A-CAGGS-Tet3G-IRES2-PuroR + pCAGGS-T2TP

pT2A-BItight-GCaMP6s-2AP-mCherry + pT2A-CAGGS-Tet3G-IRES2-PuroR + pCAGGS-T2TP

Cells were cultured as above with antibiotic replacement at 12 h and puromycin selection after 2 days.

#### PC-3 and MDA-MB-231 cells

PC-3 or MDA-MB-231 cells (1× 10^6^) were seeded into 6-well plates (1 ml of medium without PSA) for 18-24 h. A transfection mixture (4.0 μg of total plasmid DNA and 7.0-10 μl of Lipofectamine 2000 in 250 μl of OPTI-MEM) was then added. Four plasmid set was used for transfection:

pT2A-BItight-Lifeact-mCherry-TRE + pT2A-CAGGS-Tet3G-2AP-PuroR + pCAGGS-T2TP.

pT2A-BItight-Lifeact-mCherry-TRE + pT2A-CAGGS-Tet3G-2AP-PuroR

pT2A-BItight-EGFP-TRE-Orai1 E106Q + pT2A-CAGGS-Tet3G-2A-PuroR

pT2A-BItight-EGFP-TRE + pT2A-CAGGS-Tet3G-2A-PuroR.

For set 1, the medium was replaced with conventional medium containing antibiotics 12 h after transfection. After 2 days, cells were further cultured in medium containing 2-5 μg/ml puromycin for 2 days. For sets 2-4, the medium was replaced with conventional medium containing antibiotics 12 h after transfection, and cells were used for experiments one day later without puromycin selection.

### *in vitro* under-agarose assay

This method was modified from the protocol reported by Heit *et al*.^16^. To control adhesion of PGCs with dish bottom, the surface of glass bottom dish (Eppendorf) was treated by 0.5 g/L of Pluronic®F-127 (Sigma-Aldrich) for 30 min. The treated dishes was dried by air for 15 min before use.

2 x Hank’s balanced salt solution (Gibco) and FAcs medium (1mM CaCl_2_) were mixed in 1:2 and warmed up to 70℃. UltraPure^TM^ Agarose (Invitrogen) solution (48 mg/ml) was prepared by microwave. HBSS/FAcs solution and UltraPure^TM^ Agarose solution were mixed in 1.99:1.7. 3 ml of mixed solution poured onto the glass bottom and solidified at room temperature (RT). 250 μl of FAcs was added to the gel and placed at the condition of 37℃, 5% CO_2_ for 30 min for equilibration. In case of chemical addition, BAPTA (Toronto Research Chemicals Inc), 2-APB (R&D Systems), SKF96365 (Wako), or ERseeing (Funakoshi) was added to the mixed solution of HBSS/FAcs and UltraPure^TM^ Agarose and each final concentration is 100 μM, 100 μM, 50 μM, and 0.3μM, respectively.

The gel was removed from the glass bottom dish. After removing the remaining solution on the dish, 2.5×10^5^ PGCs and 5.0×10^5^ FluoSpheres^TM^ Polystyrene Microspheres (Invitrogen, F8836) with a diameter of 10 μm were suspended in 5 μl of FAcs medium and seeded on the glass bottom dish and then the gel was put back on. 50 μl of FAcs and Doxycycline (Dox; 1 μg/ml, final concentration; Clontech) were added on the gel. At this time, we adjusted the final concentration of Dox to 1 μg/ml.

### *in vitro* migration assay

PGCs (1.4×10^4^ cells/μl) and COS7 cells (2.8×10^6^ cells/μl) were suspended in 4-5% and 6-7% Matrigel^®^ Basement Membrane Matrix Growth Factor Reduced, Phenol Red Free (Corning), respectively, prepared in /FAcs medium containing 1mM CaCl_2_. The suspensions were seeded adjacent to each other in glass bottom dishes (Matsunami). The dishes were incubated at 37℃ for 20 min to allow Matrigel solidification, after which FAcs medium (1mM CaCl_2_) containing Dox (1 μg/ml, final concentration) was added on top of the Matrigel. Where indicated, inhibitors were added to FAcs medium at the following final concentrations: BAPTA (100 μM), 2-APB (100 μM), SKF96365 (50 μM).

### 3D culture of PGCs with thapsigargin or A23187 treatment

PGCs (1.4×10^4^ cells/μl) were suspended in 4-5% Matrigel, prepared by diluting in FAcs medium containing 1mM CaCl_2_, and seeded into glass bottom dishes (Matsunami). The dish was incubated at 37℃ for 20 min to allow Matrigel solidification. FAcs medium (1mM CaCl_2_) containing Dox (1 μg/ml, final concentration) was then overlaid onto the Matrigel. Thapsigargin (2 μM, final concentration) or A23187 (10 μM, final concentration) was added to the overlaid FAcs medium.

### Visualization of the endoplasmic reticulum in PGCs

STIM1-EGFP–expressing PGCs were collected by centrifugation at 1,500 rpm for 5 min and resuspended in Assay Buffer from the ER-ID^®^ Red Assay Kit (Enzo). Dual Detection Reagent and Hoechst 33342 (both included in the kit) were added at 2 μl per 1 ml of suspension and incubated for 15 min at RT. The cells were then collected again by centrifugation at 1,500 rpm for 5 min, resuspended in FAcs medium, and maintained at 37 °C with 5% CO₂ until imaging.

### PGC transplantation into embryo

For back-infusion, cultured PGCs were collected by centrifugation at 1,500 rpm for 5 minutes, washed several times in OPTI-MEM, and resuspended at 5,000 cells/μl. A 1 μl suspension was injected into the heart of HH15 chick or quail embryo using a fine glass capillary. To visualize blood vessels, Alexa Fluor™ 594- or Alexa Fluor™ 488-conjugated acetylated low-density lipoprotein (594-AcLDL or 488-AcLDL; ThermoFisher Sciencetific, L35353 and L23380) was diluted 1:2 in OPTI-MEM and co-injected with PGCs. For the tet-on induction, 1 μg/ml Dox was added to PGC cultures in FAcs medium 24 hours before transplantation. For *in ovo* induction, 0.5 ml of 100 μg/ml Dox solution was injected between the embryo and yolk as described previously^30^. Embryos were incubated at 38.5℃ until intended developmental stage.

### Transplantation of HT-1080, PC-3, and MDA-MB-231 cells into embryo

HT-1080, PC-3, and MDA-MB-231 cells were detached from culture dishes using 0.05% trypsin-EDTA in phosphate-buffered saline (PBS) and trypsinization was quenched with an equal volume of FBS. Cells were collected by centrifugation at 1,500 rpm for 5 min, washed several times with OPTI-MEM, and resuspended at 25,000 cells/μl. A 1 μl suspension was injected into the heart of HH15 chick or quail embryos using a fine glass capillary. To visualize blood vessels, either 12.5% Rhodamine-Lens culinaris lectin (Rho-LCA; Vector Laboratories, L-1040) or 12.5% FITC-conjugated Lens culinaris agglutinin (FITC-LCA; Vector Laboratories, FL-1041) diluted in OPTI-MEM was co-injected with cells. Embryos were incubated at 38.5 ℃ until the intended developmental stage.

### Fixation, whole-mount, and section immunostaining

#### Whole-mount immunostaining

Embryos containing infused PGCs were fixed in 4% PFA (paraformaldehyde) in PBS at 4 ℃ for 20 h. For immunostainings with QH1, EGFP, and mCherry in whole quail embryos, samples were washed 3 times for 60 minutes each in TNTT buffer (0.1 M Tris-HCl, pH 7.5; 0.15 M NaCl; 0.05% Tween 20; 0.5% TritonX-100) at 4 ℃. To inactivate endogenous peroxidase, embryos were immersed in 3% hydrogen peroxide in methanol for 5 days at 4 ℃, followed by 6 washes in TNTT (60 min each, 4 ℃). Blocking was performed in 1% Blocking Reagents (Roche) in TNTT for 1 hour at 4 ℃. The blocked samples were treated 4 days at 4℃ with primary antibodies diluted in blocking solution: QH1 (Hybridoma Bank, AB_531829, 1:200), anti-EGFP goat polyclonal antibody (GeneTex, GTX266673, 1:1,000), and anti-mRFP rabbit polyclonal antibody (ROCKLAND, 600-401-379, 1:1,000). After 6 washes in TNTT (60 min each, RT), samples were incubated for 4 days at 4 ℃ with secondary antibodies: Alexa Fluor 555-conjugated donkey anti-rabbit IgG (Invitrogen A31572, 1:500), HRP-linked sheep anti-mouse IgG (Cytiva NA931V, 1:500), and Alexa Fluor 488-conjugated donkey anti-goat IgG (Invitrogen A11055, 1:500) in 1% blocking solution. Finally, embryos were washed 3 times for 1 h each in TNTT (RT).

Fluorescence signal amplification was performed by incubating embryos in 1% Cyanine 5 reagent (PerkinElmer FP1171) with amplification diluent (PerkinElmer FP1135) for 15 min at RT, followed by 3 washes in TNTT (1 h each, RT). Embryos were cleared by sequential immersion in 40% glycerol/PBS (1 hour, 4 ℃), 60% glycerol/PBS (2-3 hours, 4 ℃), and 80% glycerol/DDW overnight at 4℃.

#### Section immunostaining

For section staining, fixed embryos were washed twice with PBS, and cryoprotected in 30% sucrose/PBS at 4℃ for 3-5 hrs. Samples were then incubated overnight at 4 ℃ in a 1:2 mixture of 30% sucrose/PBS and O.C.T. Compound (Sakura Finetek Japan), followed by embedding in 100% O.C.T. and freezing at -30℃. Cryosections (12 µm) were washed 3 times in TNTT and blocked in 1% Blocking Reagents/TNTT for 1 h at RT. Sections were incubated at 4℃ with primary antibodies: anti-EGFP goat polyclonal antibody (GeneTex, GTX266673, 1:1,000) and anti-mRFP rabbit polyclonal antibody (ROCKLAND, 600-401-379, 1:1,000) diluted in blocking solution. After 3 washes in TNTT (5 min each, RT), sections were incubated for 1 h at RT with secondary antibodies: Alexa Fluor 555-conjugated donkey anti-rabbit IgG (Invitrogen A31572, 1:500) and Alexa Fluor 488-conjugated donkey anti-goat IgG (Invitrogen A11055, 1:500) in 1% blocking solution. Sections were washed 3 times in TNTT and mounted in Fluoromount-G^TM^ with DAPI (Invitrogen).

### 2D culture of cancer cells with chemical inhibitors and Ca²⁺ ionophore

HT-1080, PC-3, and MDA-MB-231 cells (1.8 × 10^3^ cells/μl) were seeded onto the glass-bottom dishes (Matsunami). After incubation at 37 °C for 15 h, live imaging was initiated. The following chemicals were then added to the culture medium at their final concentrations: CK-666 (120 μM), SKF96365 (50 μM), Thapsigargin (1 μM), BAPTA (100 μM), 2-APB (75 μM), and A23187 (50 μM).

### Visualization of ER, intracellular Ca²⁺, and PM in PC-3 and MDA-MB-231 cells

PC-3 and MDA-MB-231 cells (1.8 × 10³ cells/μl) were seeded onto glass-bottom dishes (Matsunami) and incubated at 37 °C for 15 h. After three washed with PBS, cells were stained with ER-ID Dual Detection Reagent (1:1,000 dilution) for 20 min at RT, followed either by loading with Fluo-4 AM (3 μM in Opti-MEM; ThermoFisher) for 30 min at 37 °C to visualize intracellular Ca²⁺, by staining with CellMask Green (1:1,000 dilution in Opti-MEM; ThermoFisher) for 15 min at 37 °C to visualize the PM. After three additional PBS washes, cells were maintained in complete culture medium at 37 °C until imaging.

### Knockdown of IP3R3 in PC3 and MDA-MB-231 cells

PC-3 or MDA-MB-231 cells (1 × 10⁶) were seeded onto 6-well plates with 1 mL of medium without PSA and cultured for 18–24 h. The medium was then replaced with 1 mL of Opti-MEM, and the RNAi transfection mixture (containing 5 μL of DsiRNA [final concentration: 50 nM] and 5 μL of RNAiMAX (ThermoFisher Sciencetific), diluted in 100 μL of Opti-MEM) was added. Cells were incubated for 24 h, washed three times, and subsequently maintained in complete culture medium at 37 °C for an additional 72 h prior to qPCR analysis or other experiments.

DsiRNAs targeting ITPR3 (IP_3_R3) were purchased from Integrated DNA Technologies (IDT):

1. ITPR3. 13. 1: GGA CUG ACA AGA AUA ACG UGA UGC UCG UUA UUC UUG
2. ITPR3. 13. 2: GAC AUC AUG GUC ACU AAG GGU UGG GCU UAG UGA CC

### Quantitative PCR analysis of IP3R3 knockdown

Total RNA was extracted from PC-3 and MDA-MB-231 cells 72 h after transfection using the NucleoSpin RNA XS kit (Takara), according to the manufacturer’s instructions. Complementary DNA (cDNA) was synthesized using the PrimeScript™ II 1st Strand cDNA Synthesis Kit (Takara). Quantitative PCR (qPCR) was performed with SYBR™ Green qPCR Master Mix (Thermo Fisher Scientific) on a CFX Connect Real-Time PCR Detection System (Bio-Rad).

Primer sets used:

Forward: ccctcaatctgaccaacaaga

Reverse: cggtagatgaggtcaaagag

### Image acquisition and processing

#### *In ovo* live imaging

Cultured embryos was imaged using an MVX10 microscope (Olympus) or a S9 D Stereo Microscope (Leica). Images were acquired with high-speed recording software (Hamamatsu) or Flexacam C3 (Leica). Acquired images were enlarged and processes using Fiji-ImageJ (NIH).

#### *Ex vivo* live imaging

Samples on glass-bottom dishes (Matsunami) were placed in an incubation chamber (37 °C, 5% CO_2_: TOKAIHIT STX) equipped with an IX83 inverted microscope (Olympus) interfaced with a Dragonfly 200 spinning-disk confocal system (Oxford Instruments). Images were captured with a scientific camera and acquired using Fusion software (Oxford Instruments). Images were enlarged and processed using Imaris (Oxford Instruments) or Fiji-ImageJ (NIH).

#### In vitro imaging

For live imaging of under-agarose assay and *in vitro* migration assay, samples on the glass-bottom dishes (Greiner bio-one) were maintained in an incubation chamber (37 °C, 5% CO_2_: TOKAIHIT STX) on an IX83 inverted microscope with a Dragonfly 200 spinning-disk confocal system. Images were acquired with a scientific CMOS camera using Fusion software. Z-stacks were processed, enlarged, and reconstructed along the Z-axis using Imaris. ER-ID subtraction images were generated by removing ER-derived Ca^2+^ signals from Fluo-4 to highlight cytoplasmic Ca^2+^ dynamics.

#### Fixed whole-mount imaging

Whole-mount fixed samples were imaged with the Dragonfly 200 spinning-disk confocal system. Images were captured with a scientific camera and acquired using Fusion software. Acquired Z-series images were deconvoluted using Huygens (Scientific Volume Imaging) and processed for 3D reconstruction in Imaris (ver.9.5). Iso-surfaces rendering with texture mapping and plane clipping (dorsal-ventral orientation) was performed for visualization.

### Quantification and statistical analysis

#### PGC quantification

Areas and fluorescence intensities of GCaMP6s and mCherry signals in blebs and cell bodies (*in vitro* under-agarose assay and *in vivo*) were measured using ImageJ. Migration tracks and distances in the *in vitro* migration assay were quantified using the Manual tracking plug-in in imageJ. The number of protrusion-forming PGCs (*in vitro* migration assay and Ex-VaP) and arrested PGCs (5 h after infusion in Ex-VaP) were counted manually. EGFP+ or mCherry+ PGCs were also counted manually at 6 h after infusion in Ex-VaP and at 2 days after infusion in the dorsal mesentery and gonads.

#### Cancer cell quantification

Protrusion formation in 2D culture was quantified manually. The number of control or Orai1 E106Q-expressing HT-1080 cells passing through the imaging field in the YSV was manually counted from time-lapse movies. Extravasated cancer cells were quantified by counting EGFP+ cells in the YSV at 3.5 h after infusion.

All graphs were generated using Microsoft Excel. Statistical comparisons were performed using a two-tailed, unpaired Student’s t-test. In box plots, the box represents the interquartile range (IQR), the central line indicates the median, and the mean is marked by a cross (×). Whiskers represent the standard error of the mean (s.e.m.), unless otherwise specified in the figure legends.

#### RNA-seq data analysis

To assess the expression of STIM1/2 and ORAI1/2 transcripts, we analyzed published RNA-seq data of chicken PGCs (GSE188689). Transcript abundance was quantified as transcripts per million (TPM) using *kallisto* (v0.46.0) with default settings, pseudoaligning the reads to the chicken reference transcriptome (GCF_016699485.2).

## FIGURE LEGENDS

**Extended Data Figure 1.**

**Endogenous PGCs form membrane blebs during TEM.**

**a**, F-actin distribution in a chick endogenous PGC undergoing TEM within the Ex-VaP. White dotted lines delineate the boundary between the intravascular space (IS) and the extravascular space (ES); yellow dotted line outlines the cell. A membrane protrusion lacking cortical actin is marked by yellow arrowheads, indicative of a bleb. Nucleus stained with DAPI (blue). Scale bar, 10μm.

**Extended Data Figure 2.**

**Ca^2+^ influx and actomyosin Dynamics during bleb formation in chick PGCs.**

**a,** Schematic of the under-agarose compression (UAC) assay used to apply mild physical constraint to PGCs, promoting bleb formation. **b,** Representative image of a PGC expressing mCheery, showing multiple blebs (white arrowheads). Cell outline is marked with a white dotted line. **c,** Time-lapse images of a PGC expressing MRLC-EGFP showing a typical bleb cycle. **d,** Time-lapse images (Supplementary Movie 5, 6) of a PGC co-expressing EGFP-CAAX and Lifeact-mCheery. Yellow and white arrowheads denote extending and retracting blebs, respectively. **e,** Time-lapse images (Supplementary Movie 7) of PGC expressing GCaMP6s, highlighting a bleb region (white arrowheads). **f,** Changes in GCaMP6s fluorescence in the bleb and cell body during a bleb cycle. *N*=3 cells per region. Intensities are normalized to co-exressed cytoplasmic mCherry. E, extension phase; R, retraction phase. **g,** Ratio of GCaMP6s fluorescence between bleb and cell body under UAC assay. *N*=10. ***p < 0.001 (two-sided paired Student’s *t*-test). Error bars, s.e.m. Scale bars, 2.5 μm (**b, d**), 3 μm (**c, e**).

**Extended Data Figure 3.**

**Subcellular localization of STIM1 and Orai1 in chick PGCs.**

**a,** Confocal images of PGC expressing STIM1-EGFP (green), counterstained with ER-ID (red) and Hoechst (blue), showing localization of STIM1 to the ER. **b,** Co-expression of Stim1-EGFP (green) and Orai1-mCherry (red) in a PGC reveals the distinct spatial localization of STIM1 in the ER and Orai1 at the PM, enabling visualization of their relative positions under resting conditions. Scale bars, 5 μm.

**Extended Data Figure 4.**

**Ca^2+^ ionophore triggers bleb formation in PGCs and human cancer cells.**

**a-d**, Time-lapse imaging of a PGC (a) and HT-1080 (b), PC-3 (c), MDA-MB-231 (d) cell exposed to the Ca^2+^ ionophore A23187. Time is indicated in seconds relative to the initial addition of A23187 (0s). Scale bars, 5 μm.

**Extended Data Figure 5.**

**SCF enhances chick PGC migration in a 3D co-culture assay.**

**a,** Schematic and representative images of the Matrigel-based co-culture system using GCaMP6s^+^ PGCs (green) and COS7 cells expressing Gap-TdTomato (red). **b,** Time-lapse images of PGCs under three conditions: co-culture with COS7 cells expressing Gap-TdTomato alone (control), with SDF-1, or with SCF. Arrowheads of different colors mark individual PGCs; with corresponding dotted lines traceing their trajectories. **c,** Migration trajectories of PGCs co-cultured with COS7 cells expressing Gap-TdTomato alone (control) or with SCF (COS7 cells positioned to the left; *N*=15 cells). **d**, Migration trajectories of PGCs co-cultured with SCF in the presence of SKF96365 (*N*=15). **e,** Migration distances of PGCs over 30 minutes (*N*=15). **f,** Proportion of PGCs forming protrusions (*N*=30). **g**, Time-lapse images of a migrating PGC showing sustained frontal blebbing under SCF stimulation. White dotted lines mark the initial cell contour, indicating net displacement. **h**, Quantification of GCaMP6s fluorescence intensity ratio between front and rear regions of migrating PGCs (*N*=13 cells). **i,** Confocal images of PGCs co-expressing Stim1-EGFP (green) and Orai1-mCheery (red), showing colocalization at ER-PM contact sites prior to bleb formation (white arrowheads). **j**, Percentage of PGCs exhibiting blebs under SCF stimulation with or without SOCE inhibition by SKF96365 (*N*=3). Error bars, s.e.m. *P<0.05, **P<0.01, ***P<0.001. (two-sided unpaired Student’s *t*-test). Scale bars, 30 μm (**a**), 15 μm (**b**), 5 μm (**g**), 1 μm (**i**).

**Extended Data Figure 6.**

**Orai1 E106Q inhibits bleb formation and migration in chick PGCs.**

**a,** Representative images of PGCs expressing EGFP or Orai1 E106Q under the UAC assay. White dotted lines mark cell boundaries. **b,** Quantification of the proportion of total cell area occupied by bleb regions under the UAC assay. **c,** Representative migration trajectories of PGCs expressing EGFP or Orai1 E106Q under SCF stimulation. **d**, Migration distances over 30 minutes in the SCF assay. *N*=12 cells per condition. *P<0.05, ***P<0.001 (Two-sided, unpaired Student’s *t*-test). Scale bars, 5 μm.

**Extended Data Figure 7. SOCE is essential for chick PGC migration *in vivo*.**

**a,** Schematic of the experimental timeline: EGFP or Orai1 E106Q-expressing PGCs into the vasculature of HH15 chick embryos, together with an equal number of mCheery^+^ PGCs as internal infusion control. PGCs were analyzed for vascular arrest at the Ex-VaP and subsequent tissue migration to the dorsal mesentery and gonads. **b,** Representative fluorescence images of the Ex-VaP region at 5 hpt (HH16), showing comparable vascular arrest of mCheery^+^ and Orai1 E106Q^+^ (EGFP^+^) PGCs. **c,** Quantification of intravascular arrest, calculate as the ratio of Orai1 E106Q^+^ PGCs to co-infused mCherry^+^ PGCs (*N*=5 embryos). **d,** Quantification of PGC migration distance over 60 minutes for Orai1 E106Q^+^and control PGCs (*N*=8 cells, respectively). **e,** Horizontal sections of E4.5 chick embryos (2 days post-transplantation) showing localization of EGFP^+^ (control), Orai1 E106Q^+^ (EGFP^+^), and co-infused mCherry^+^ PGCs in the dorsal mesentery (DM, white dotted line) and gonadal primordium (G, yellow dotted line). **f,** Quantification of PGC localization at E4.5. Ratios of EGFP^+^ or Orai1 E106Q^+^ cells in the DM and G were normalized to the number of co-infused mCherry^+^ PGCs (*N*=4 embryos). **g,** Cell viability over time for mCherry^+^ control (blue) and Orai1 E106Q^+^ (red) PGCs. *N*=5. **h,** Proliferation of individual PGCs (*N*=26 for control, *N*=29 for Orai1 E106Q^+^) following Dox-induced expression at Day0. Error bars, s.e.m. “X” indicates the mean. **P<0.01, ***P<0.001 (two-sided unpaired Student’s *t*-test). Scale bars, 1000 μm (**b**), 100 μm (**e**).

**Extended Data Figure 8.**

**SOCE drives bleb-based TEM of HT-1080 cells.**

**a,** Schematic of the infusion strategy introducing HT-1080 cells into the vasculature of HH15 chick embryos. **b,** Representative images of an HT-1080 cell undergoing TEM in the YSV. Lifeact-mCherry reveals blebs (actin-poor; yellow dotted line) and actin-rich protrusions (White arrowheads). White dotted lines delineate intravascular (IS) and extravascular (ES) compartments. **c,** Time-lapse imaging of an HT-1080 cell exposed to thapsigargin in 2D culture. Time is shown in seconds relative to the initial addition. The yellow dotted lines outline the emerging blebs. **d,** Quantification of the number of blebs in thapsigargin treated HT-1080 cells in 2D culture. (*N*=7 cells). **e,** Control or Orai1 E106Q-expressing HT-1080 cells (green) in the lateral YSV region at 20 mpt. Vasculature was visualized with Rho-LCA (red). **f,** Quantification of control or Orai1 E106Q-expressing HT-1080 cells passing through the YSV imaging field at 3–4 and 23-24 mpt (*N*=6 embryos). **g,** Control and Orai1 E106Q-expressing HT-1080 cells in the lateral YSV at 1.5 hpt. Arrowheads indicate actin-rich protrusions. **h,** Quantification of the proportion of cells forming actin-rich protrusions at 1.5 hpt (*N*=5 embryos). **i,** Transmigrating control and Orai1 E106Q+ HT-1080 cells (green) in the YSV at 3.5 hpt. Vasculature was visualized with Rho-LCA (red). ***P<0.001 (**d**: two-sided paired Student’s *t-*test; **f**, **h**: two-sided unpaired Student’s *t*-test). Scale bars: 12 μm in **b**, 5 μm in **c,** 10 μm in **e, g,** 30 μm in **i**.

**Extended Data Figure 9.**

**Bleb formation and TEM of epithelial cancer cells are SOCE-independent.**

**a, b,** Time-lapse imaging of PC-3 (**a**) and MDA-MB-231 (**b**) cells exposed to SKF96365 in 2D culture. Time is shown in seconds relative to the initial addition of SKF96365. Yellow dotted lines outline the blebs. **c,** Quantification of bleb numbers before and after SKF96365 treatment in PC-3 and MDA-MB-231 (*N*=8 and 6 cells, respectively). **d, e,** Time-lapse imaging of PC-3 (**d**) and MDA-MB-231 (**e**) cells exposed to thapsigargin in 2D culture. **f,** 3D reconstructions of control and Orai1 E106Q-expressing PC-3 or MDA-MB-231 cells (green iso-surfaces) undergoing TEM in the YSV at 3.5 hpt. Blood vessels were labeled by intravenous injection of Rho-LCA (brown). Endothelial volumes were digitally clipped to visualize transmigrating cells. **g,** Images of transmigrated control or Orai1 E106Q-expressing PC-3 or MDA-MB-231 cell (green) in the YSV at 3.5 hpt. Blood vessels were visualized by injection of Rho-LCA (red). White dotted lines delineate intravascular (IS) and extravascular (ES) compartments. **h,** Quantification of extravasation efficiency for control and Orai1 E106Q-expressing PC-3 and MDA-MB-231 cells. Efficiency was calculated as the percentage of EGFP+ or Orai1 E106Q-expressing cells located outside the vasculature among all labeled cells in the YSV (*N* = 4 embryos per group). **i, j,** Time-lapse imaging of PC-3 (**i**) and MDA-MB-231 (**j**) cells exposed to BAPTA in 2D culture. **k,** Quantification of the number of blebs before and after BAPTA treatment in PC-3 and MDA-MB-231 (*N*=8 and 6 cells, respectively). Error bars, s.e.m. ns, not significant (**c, k**: two-sided paired Student’s *t-*test; **h**: Two-sided Welch’s t-test). Scale bars: 5 μm in **a**, **b, d, e, i, j, n,** 20 μm in **f, g**, 2 μm in **i, j.**

**Extended Data Figure 10.**

**IP_3_R is necessary for bleb formation and TEM of epithelial cancer cells.**

**a**, Dual imaging of ER-ID and Fluo-4 in a PC-3 cell during bleb formation. Left and middle: raw images; right: subtraction of ER–ID signal from Fluo-4. **b,** Dual visualization of CellMask (green) and ER-ID (red) showing ER tubules extending into a bleb of a PC-3 cell (white arrowhead). **c,** Time-lapse imaging of an MDA-MB-231 cell before and after 2-APB addition in 2D culture. Time is shown in seconds relative to the initial addition of 2-APB. The yellow dotted lines outline the emerging blebs. **d,** Knockdown efficiency of three different DsiRNAs targeting IP_3_R3 in PC-3 and MDA-MB-231 cells. Relative expression levels were normalized to IP_3_R3 expression in the negative control (set to 1). β-actin was used as a housekeeping gene. **e**, Representative images of Lifeact-expressing PC-3 cells: control, IP_3_R3 KD1, and KD2, in 2D culture. **f,** Quantification of the proportion of bleb-forming PC-3 cells in 2D culture. **g,** Quantification of the proportion of PC-3 cells with actin-rich protrusions in 2D culture. **h,** Representative images of control, IP_3_R3 KD1, and KD2 PC-3 and MDA-MB-231 cells in the YSV at 20 mpt. Blood vessels were visualized by injection of Rho-LCA (red). White dotted lines delineate the boundary between the intravascular space (IS) and the extravascular space (ES). **i, j,** Quantification of control, IP_3_R3 KD1, and KD2 cells in PC-3 (**i;** *N*=6, 6, 4 embryos, respectively) and MDA-MB-231 (**j;** *N*=4 embryos, each) during passage through the YSV imaging field at 3–4 and 23-24 mpt. **k**, Representative images of control, IP_3_R3 KD1, and KD2 PC-3 and MDA-MB-231 cells in the YSV at 3.5 hpt. **l, m**, 3D reconstructions of IP_3_R3 KD2 PC-3 cells (**l**) and control, IP_3_R3 KD1, and KD2 MDA-MB-231 cells (**m**) (green iso-surfaces) undergoing TEM in the YSV at 3.5 hpt. Blood vessels were labeled by intravenous injection of Rho-LCA (brown). Endothelial volumes were digitally clipped to visualize transmigrating cells. ns, not significant. *P<0.05, **P<0.01, ***P<0.001 (**d, f, g**: Two-sided Welch’s t-test; **i, j**: two-sided, un-paired Student’s *t-*test). Error bars represent standard deviation. Scale bars: 2 μm in **a-b,** 5 μm in **c, e** 10 μm in **h**, **k, l, m.**

## VIDEO CAPTIONS

**Supplementary Videos1 |** Bleb formation during intravascular crawling of an EGFP^+^ PGC in the Ex-VaP (related to Fig. 1a). Multiple small, transient blebs (< 5 min) emerge in random directions. Time-lapse from an HH15 chick embryo (∼23 min total), acquired with a scientific CMOS camera and Fusion software. Blood vessels labeled with 594-AcLDL.

**Supplementary Videos2 |** Bleb formation during transendothelial migration (TEM) of an EGFP^+^ PGC in the Ex-VaP (related to Fig. 1b). A single, large, persistent bleb (15–30 min) emerges toward the extravascular space. Recorded from an HH15 chick embryo (∼52 min total).

**Supplementary Videos3 |** Bleb formation of Lifeact–mCherry-expressing PGCs in the UA assay (Extended Data Fig. 2d). ∼2 min 52 s acquisition.

**Supplementary Videos4 |** Bleb formation of GCaMP6s-expressing PGCs in the UA assay (Extended Data Fig. 2e). ∼13 min 51 s acquisition.

**Supplementary Videos5 |** mCherry-expressing PGCs following A23187 exposure (Extended Data Fig. 4a). ∼8 min 40 s acquisition.

**Supplementary Videos6 |** Intravascular Orai1 E106Q-expressing PGCs in the Ex-VaP (Fig. 3b). HH15 chicken embryo, ∼54 min 30 s acquisition. Blood vessels labeled with 594-AcLDL.

**Supplementary Video 7** | Lifeact–mCherry-expressing HT-1080 cells following CK-666 and SKF96365 treatment (Fig. 4d). ∼19 min 20 s acquisition.

**Supplementary Video 8** | ZsGreen1-expressing HT-1080 cells following A23187 exposure (Extended Data Fig. 4b). ∼6 min 40 s acquisition.

**Supplementary Video 9** | EGFP-expressing PC-3 cells following A23187 exposure (Extended Data Fig. 4c). ∼6 min 20 s acquisition.

**Supplementary Video 10** | EGFP-expressing MDA-MB-231 cells following A23187 exposure (Extended Data Fig. 4d). ∼5 min 40 s acquisition.

**Supplementary Video 11** | Lifeact–mCherry-expressing PC-3 cells following SKF96365 treatment (Extended Data Fig. 9a). ∼8 min acquisition.

**Supplementary Video 12** | Lifeact–mCherry-expressing MDA-MB-231 cells following SKF96365 treatment (Extended Data Fig. 9b). ∼11 min 20 s acquisition.

**Supplementary Video 13** | Lifeact–mCherry-expressing PC-3 cells following BAPTA treatment (Extended Data Fig. 9i). ∼15 min acquisition.

**Supplementary Video 14** | Lifeact–mCherry-expressing MDA-MB-231 cells following BAPTA treatment (Extended Data Fig. 9j). ∼11 min 20 s acquisition.

**Supplementary Video 15** | Lifeact–mCherry-expressing PC-3 cells following 2-APB treatment (Fig. 5f). ∼17 min 40 s acquisition.

**Supplementary Video 16** | Lifeact–mCherry-expressing MDA-MB-231 cells following 2-APB treatment (Extended Data Fig. 10c). ∼17 min 20 s acquisition.

## ACKNOWLEDGEMENTS

For 3D image processing, we thank the Center for Advanced Instrumental and Educational Support of the Faculty of Agriculture, Kyushu University. This work was supported by the following grants: JSPS KAKENHI (Grant number 22H02634 for D. S., and JP 19H04775 for Y. A.) and Princess Takamatsu Cancer Research Fund, Shinnihon Foundation of Advanced Medical Treatment Research, Terumo Life Science Foundation for D. S.

## AUTHOR CONTRIBUTIONS

M. Morita and D. S. designed the study. M. Morita performed most experiments and analyzed data. M. Morimoto performed the immunostaining experiments. T. T. performed 3D image acquisition. Y. H. and B. P. analyzed RNA-seq data. D. S., T. T., J. I., and Y. A. supervised the study. M. Morita and D. S. wrote the manuscript.

