## Extended Data 1-10 for "Bleb-based extravasation: conserved morphodynamics, divergent calcium control"

**a**

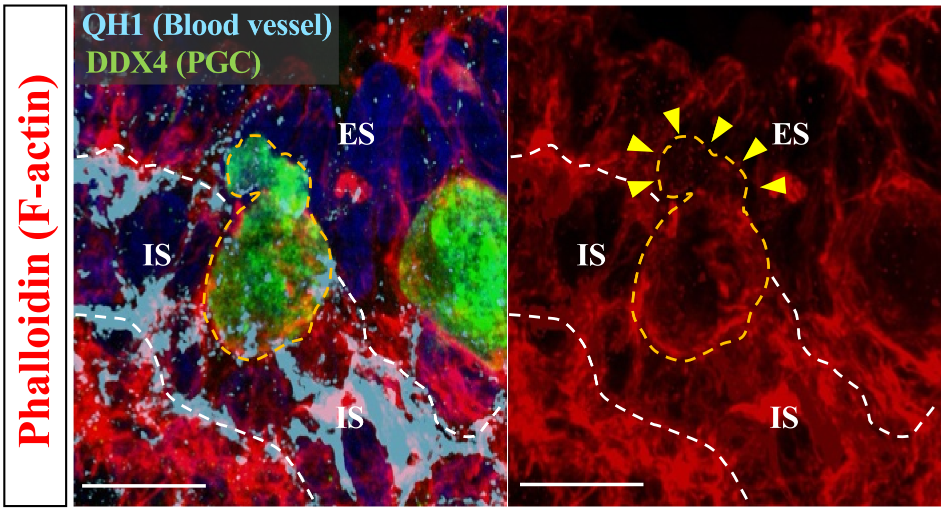


**Extended Data Figure 1 Morita *et al.***

**Extended Data Figure 1.**

**Endogenous PGCs form membrane blebs during TEM.**

**a**, F-actin distribution in a chick endogenous PGC undergoing TEM within the Ex-VaP. White dotted lines delineate the boundary between the intravascular space (IS) and the extravascular space (ES); yellow dotted line outlines the cell. A membrane protrusion lacking cortical actin is marked by yellow arrowheads, indicative of a bleb. Nucleus stained with DAPI (blue). Scale bar, 10μm.

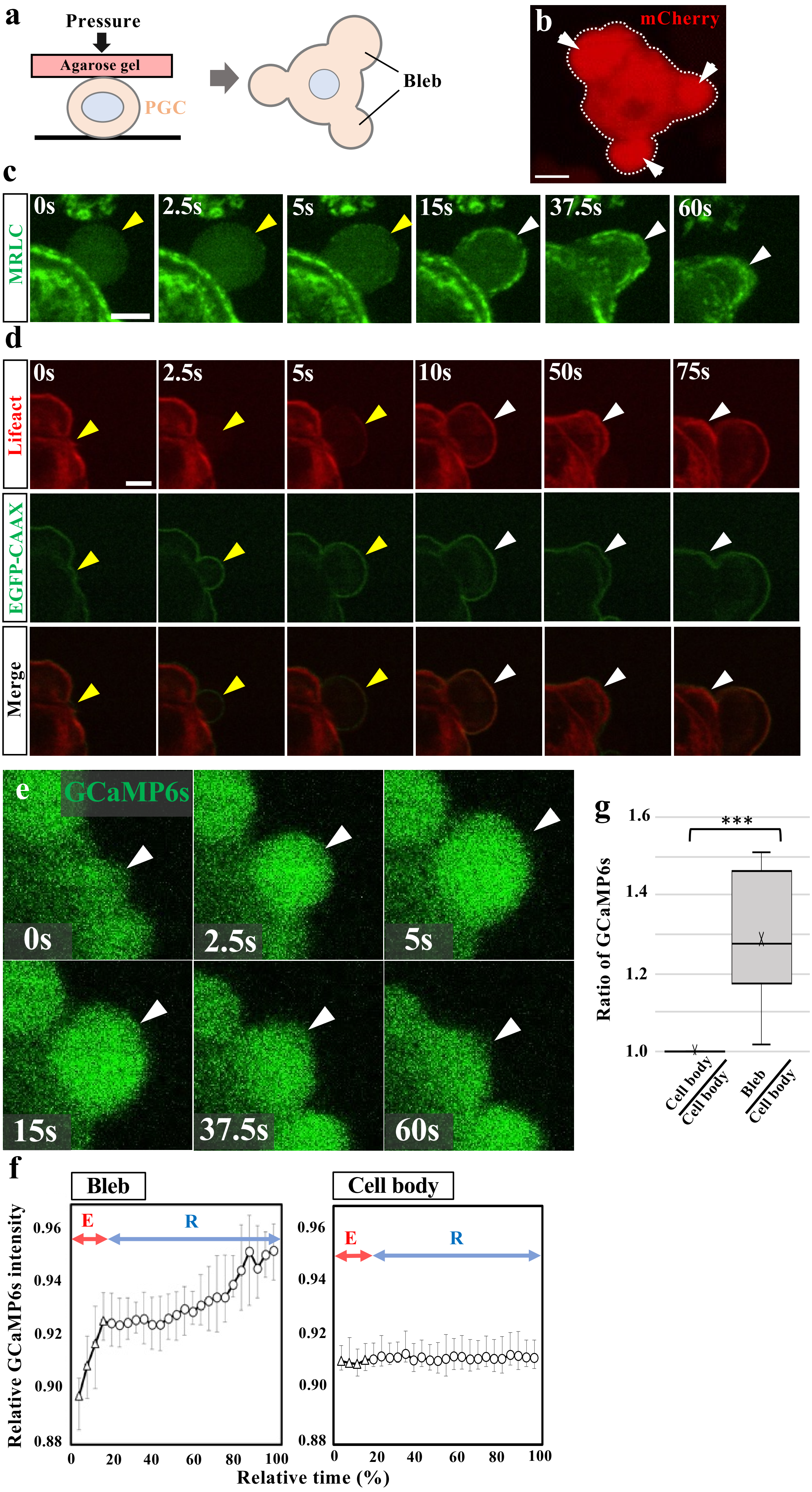


Extended Data Figure 2 Morita *et al.*

**Extended Data Figure 2.**

**$Ca^{2+}$ influx and actomyosin Dynamics during bleb formation in chick PGCs.**

**a,** Schematic of the under-agarose compression (UAC) assay used to apply mild physical constraint to PGCs, promoting bleb formation. **b,** Representative image of a PGC expressing mCheery, showing multiple blebs (white arrowheads). Cell outline is marked with a white dotted line. **c,** Time-lapse images of a PGC expressing MRLC-EGFP showing a typical bleb cycle. **d,** Time-lapse images (Supplementary Movie 5, 6) of a PGC co-expressing EGFP-CAAX and Lifeact-mCheery. Yellow and white arrowheads denote extending and retracting blebs, respectively. **e,** Time-lapse images (Supplementary Movie 7) of PGC expressing GCaMP6s, highlighting a bleb region (white arrowheads). **f,** Changes in GCaMP6s fluorescence in the bleb and cell body during a bleb cycle. $N$=3 cells per region. Intensities are normalized to co-exressed cytoplasmic mCherry. E, extension phase; R, retraction phase. **g,** Ratio of GCaMP6s fluorescence between bleb and cell body under UAC assay. $N$=10. ***$p < 0.001$ (two-sided paired Student's $t$-test). Error bars, s.e.m. Scale bars, 2.5 μm (**b, d**), 3 μm (**c, e**).

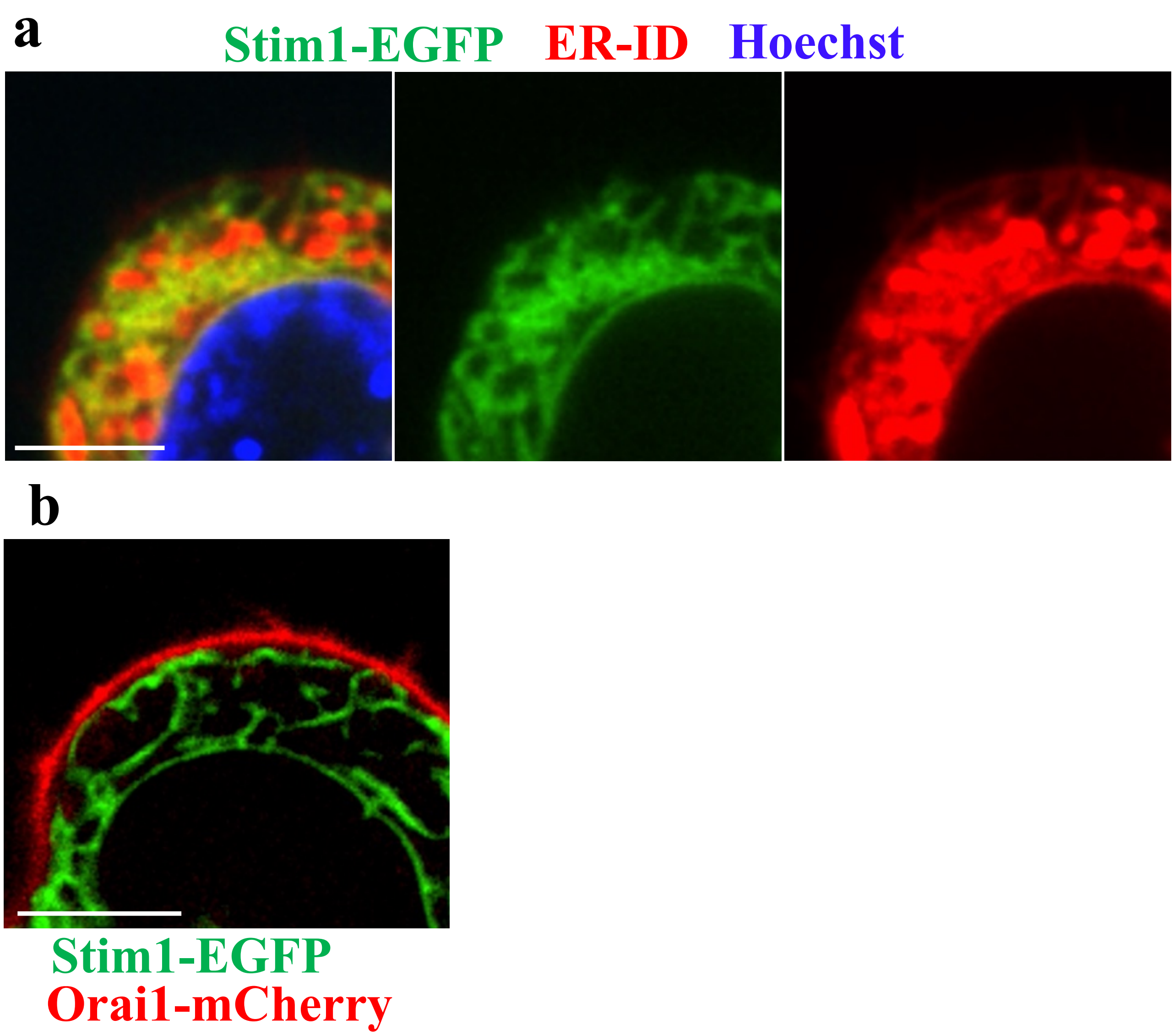


**Extended Data Figure 3 Morita *et al.***

**Extended Data Figure 3.**
**Subcellular localization of STIM1 and Orai1 in chick PGCs.**
**a,** Confocal images of PGC expressing STIM1-EGFP (green), counterstained with ER-ID (red) and Hoechst (blue), showing localization of STIM1 to the ER. **b,** Co-expression of Stim1-EGFP (green) and Orai1-mCherry (red) in a PGC reveals the distinct spatial localization of STIM1 in the ER and Orai1 at the PM, enabling visualization of their relative positions under resting conditions. Scale bars, 5 µm.

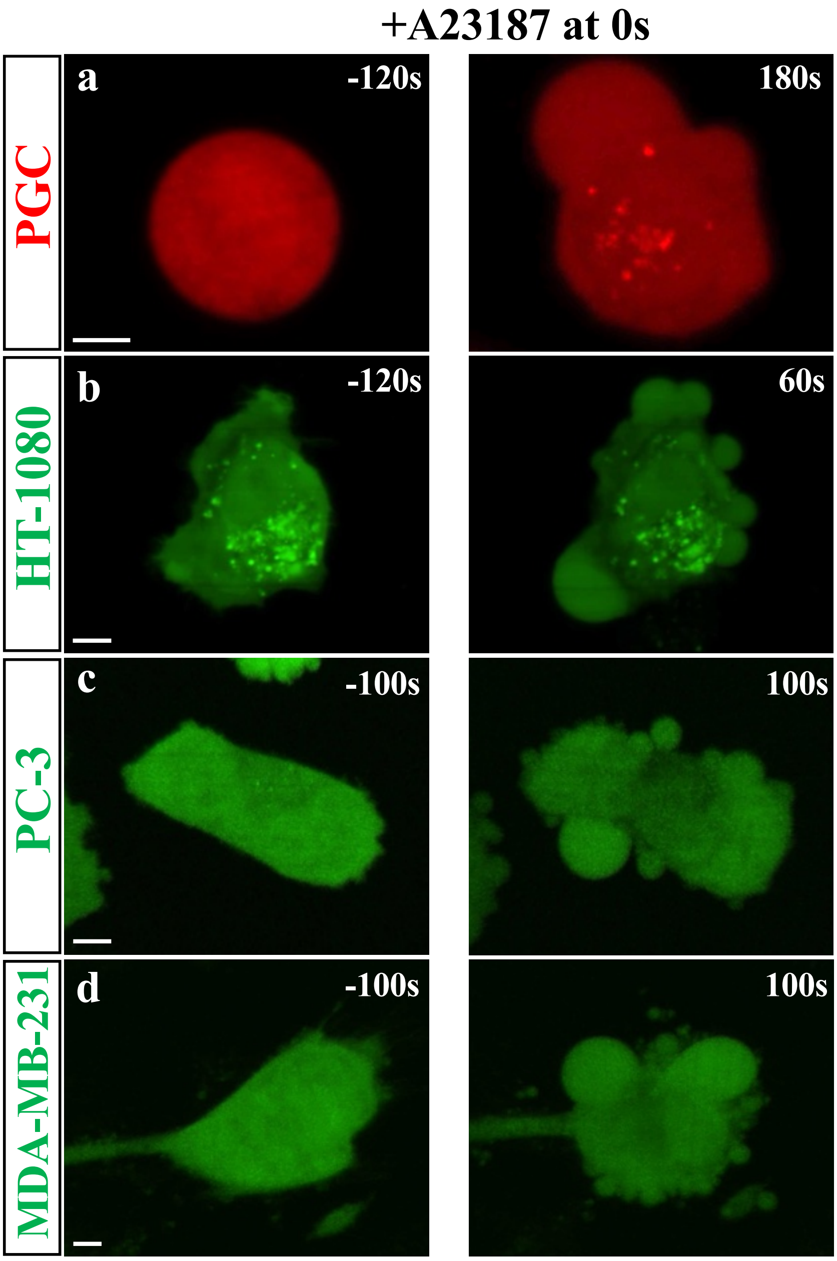


**Extended Data Figure 4 Morita *et al.***

**Extended Data Figure 4.**
**$Ca^{2+}$ ionophore triggers bleb formation in PGCs and human cancer cells.**
**a-d**, Time-lapse imaging of a PGC (a) and HT-1080 (b), PC-3 (c), MDA-MB-231 (d) cell exposed to the $Ca^{2+}$ ionophore A23187. Time is indicated in seconds relative to the initial addition of A23187 (0s). Scale bars, 5 µm.

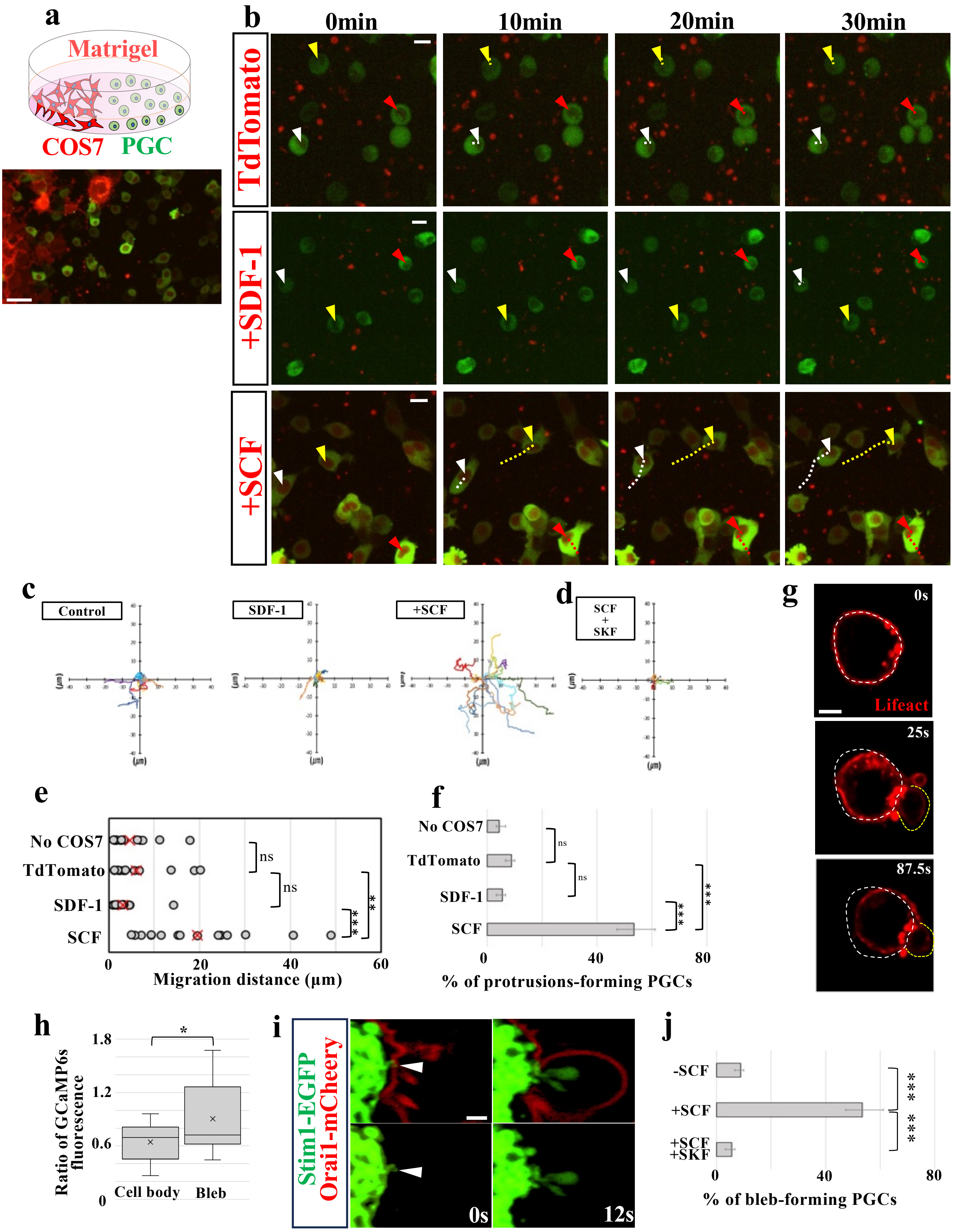


Extended Data Figure 5 Morita *et al.*

**Extended Data Figure 5.**

**SCF enhances chick PGC migration in a 3D co-culture assay.**

**a,** Schematic and representative images of the Matrigel-based co-culture system using GCaMP6s$^+$ PGCs (green) and COS7 cells expressing Gap-TdTomato (red). **b,** Time-lapse images of PGCs under three conditions: co-culture with COS7 cells expressing Gap-TdTomato alone (control), with SDF-1, or with SCF. Arrowheads of different colors mark individual PGCs; with corresponding dotted lines traceing their trajectories. **c,** Migration trajectories of PGCs co-cultured with COS7 cells expressing Gap-TdTomato alone (control) or with SCF (COS7 cells positioned to the left; *N*=15 cells). **d**, Migration trajectories of PGCs co-cultured with SCF in the presence of SKF96365 (*N*=15). **e,** Migration distances of PGCs over 30 minutes (*N*=15). **f,** Proportion of PGCs forming protrusions (*N*=30). **g**, Time-lapse images of a migrating PGC showing sustained frontal blebbing under SCF stimulation. White dotted lines mark the initial cell contour, indicating net displacement. **h**, Quantification of GCaMP6s fluorescence intensity ratio between front and rear regions of migrating PGCs (*N*=13 cells). **i,** Confocal images of PGCs co-expressing Stim1-EGFP (green) and Orai1-mCheery (red), showing colocalization at ER-PM contact sites prior to bleb formation (white arrowheads). **j**, Percentage of PGCs exhibiting blebs under SCF stimulation with or without SOCE inhibition by SKF96365 (*N*=3). Error bars, s.e.m. *P<0.05, **P<0.01, ***P<0.001. (two-sided unpaired Student's *t*-test). Scale bars, 30 μm (**a**), 15 μm (**b**), 5 μm (**g**), 1 μm (**i**).

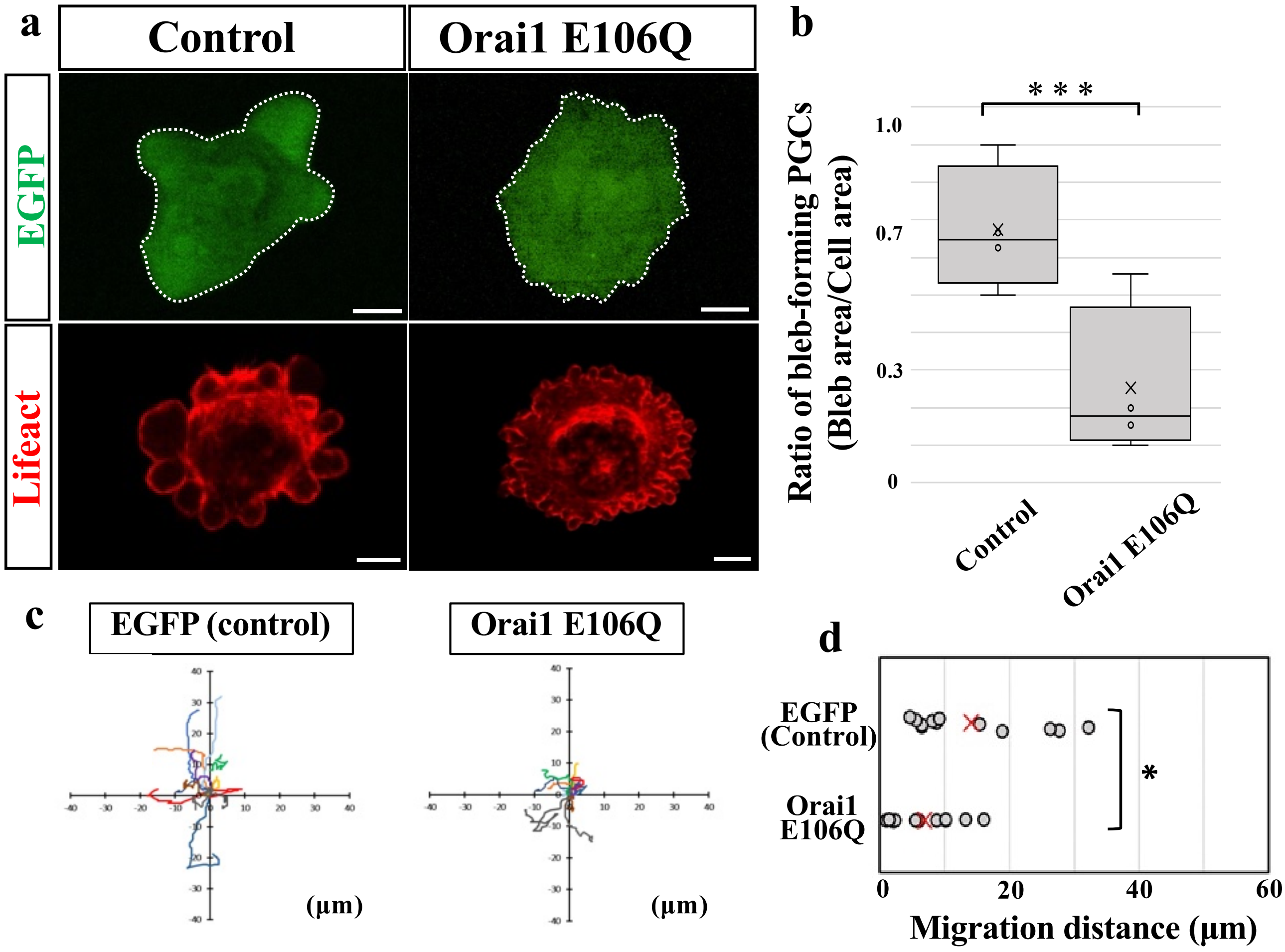


Extended Data Figure 6 Morita *et al.*

**Extended Data Figure 6.**
**Orai1 E106Q inhibits bleb formation and migration in chick PGCs.**
**a,** Representative images of PGCs expressing EGFP or Orai1 E106Q under the UAC assay. White dotted lines mark cell boundaries. **b,** Quantification of the proportion of total cell area occupied by bleb regions under the UAC assay. **c,** Representative migration trajectories of PGCs expressing EGFP or Orai1 E106Q under SCF stimulation. **d**, Migration distances over 30 minutes in the SCF assay. *N*=12 cells per condition. *P<0.05, ***P<0.001 (Two-sided, unpaired Student's *t*-test). Scale bars, 5 µm.

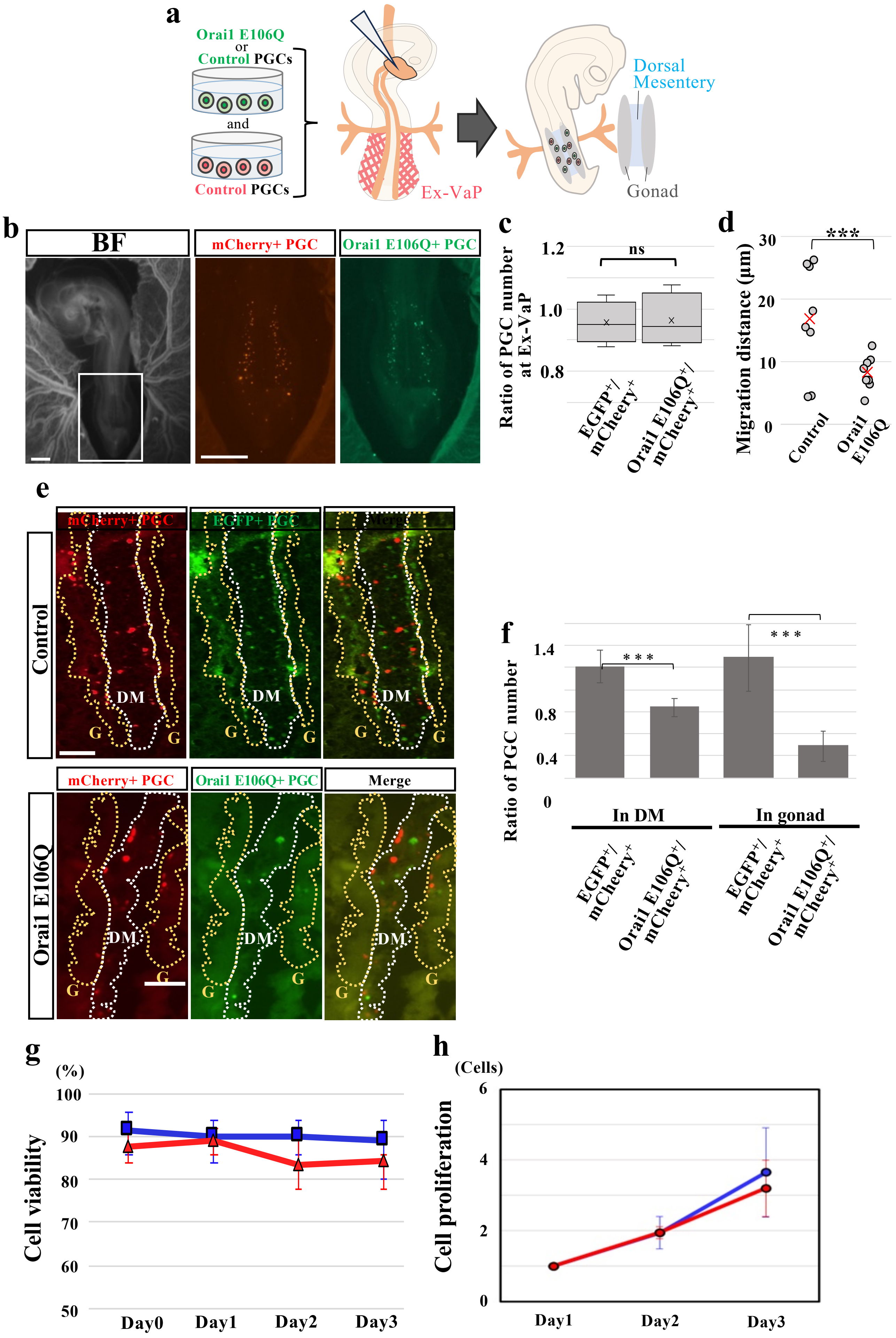


**Extended Data Figure 7 Morita *et al.***

**Extended Data Figure 7. SOCE is essential for chick PGC migration *in vivo.***

**a,** Schematic of the experimental timeline: EGFP or Orai1 E106Q-expressing PGCs into the vasculature of HH15 chick embryos, together with an equal number of $mCheery^+$ PGCs as internal infusion control. PGCs were analyzed for vascular arrest at the Ex-VaP and subsequent tissue migration to the dorsal mesentery and gonads. **b,** Representative fluorescence images of the Ex-VaP region at 5 hpt (HH16), showing comparable vascular arrest of $mCheery^+$ and Orai1 $E106Q^+$ ($EGFP^+$) PGCs. **c,** Quantification of intravascular arrest, calculate as the ratio of Orai1 $E106Q^+$ PGCs to co-infused $mCherry^+$ PGCs ($N$=5 embryos). **d,** Quantification of PGC migration distance over 60 minutes for Orai1 $E106Q^+$ and control PGCs ($N$=8 cells, respectively). **e,** Horizontal sections of E4.5 chick embryos (2 days post-transplantation) showing localization of $EGFP^+$ (control), Orai1 $E106Q^+$ ($EGFP^+$), and co-infused $mCherry^+$ PGCs in the dorsal mesentery (DM, white dotted line) and gonadal primordium (G, yellow dotted line). **f,** Quantification of PGC localization at E4.5. Ratios of $EGFP^+$ or Orai1 $E106Q^+$ cells in the DM and G were normalized to the number of co-infused $mCherry^+$ PGCs ($N$=4 embryos). **g,** Cell viability over time for $mCherry^+$ control (blue) and Orai1 $E106Q^+$ (red) PGCs. $N$=5. **h,** Proliferation of individual PGCs ($N$=26 for control, $N$=29 for Orai1 $E106Q^+$) following Dox-induced expression at Day0. Error bars, s.e.m. “X” indicates the mean. **P<0.01, ***P<0.001 (two-sided unpaired Student’s $t$-test). Scale bars, 1000 μm (**b**), 100 μm (**e**).

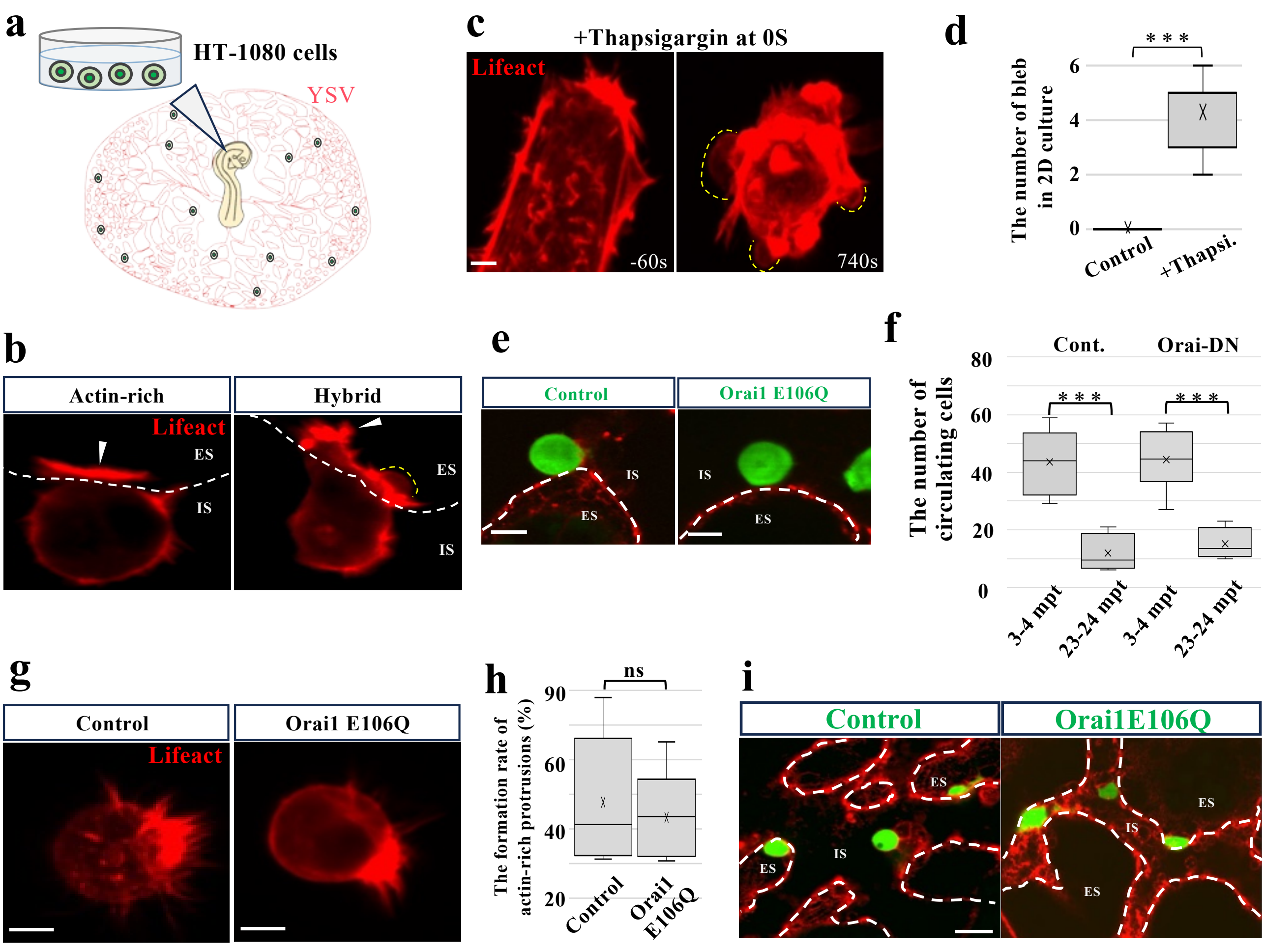


**Extended Data Figure 8 Morita *et al.***

**Extended Data Figure 8.**

**SOCE drives bleb-based TEM of HT-1080 cells.**

**a,** Schematic of the infusion strategy introducing HT-1080 cells into the vasculature of HH15 chick embryos. **b,** Representative images of an HT-1080 cell undergoing TEM in the YSV. Lifeact-mCherry reveals blebs (actin-poor; yellow dotted line) and actin-rich protrusions (White arrowheads). White dotted lines delineate intravascular (IS) and extravascular (ES) compartments. **c,** Time-lapse imaging of an HT-1080 cell exposed to thapsigargin in 2D culture. Time is shown in seconds relative to the initial addition. The yellow dotted lines outline the emerging blebs. **d,** Quantification of the number of blebs in thapsigargin treated HT-1080 cells in 2D culture. (*N*=7 cells). **e,** Control or Orai1 E106Q-expressing HT-1080 cells (green) in the lateral YSV region at 20 mpt. Vasculature was visualized with Rho-LCA (red). **f,** Quantification of control or Orai1 E106Q-expressing HT-1080 cells passing through the YSV imaging field at 3–4 and 23-24 mpt (*N*=6 embryos). **g,** Control and Orai1 E106Q-expressing HT-1080 cells in the lateral YSV at 1.5 hpt. Arrowheads indicate actin-rich protrusions. **h,** Quantification of the proportion of cells forming actin-rich protrusions at 1.5 hpt (*N*=5 embryos). **i,** Transmigrating control and Orai1 E106Q+ HT-1080 cells (green) in the YSV at 3.5 hpt. Vasculature was visualized with Rho-LCA (red). ***P<0.001 (**d**: two-sided paired Student's *t*-test; **f**, **h**: two-sided unpaired Student's *t*-test). Scale bars: 12 μm in **b**, 5 μm in **c,** 10 μm in **e, g,** 30 μm in **i**.

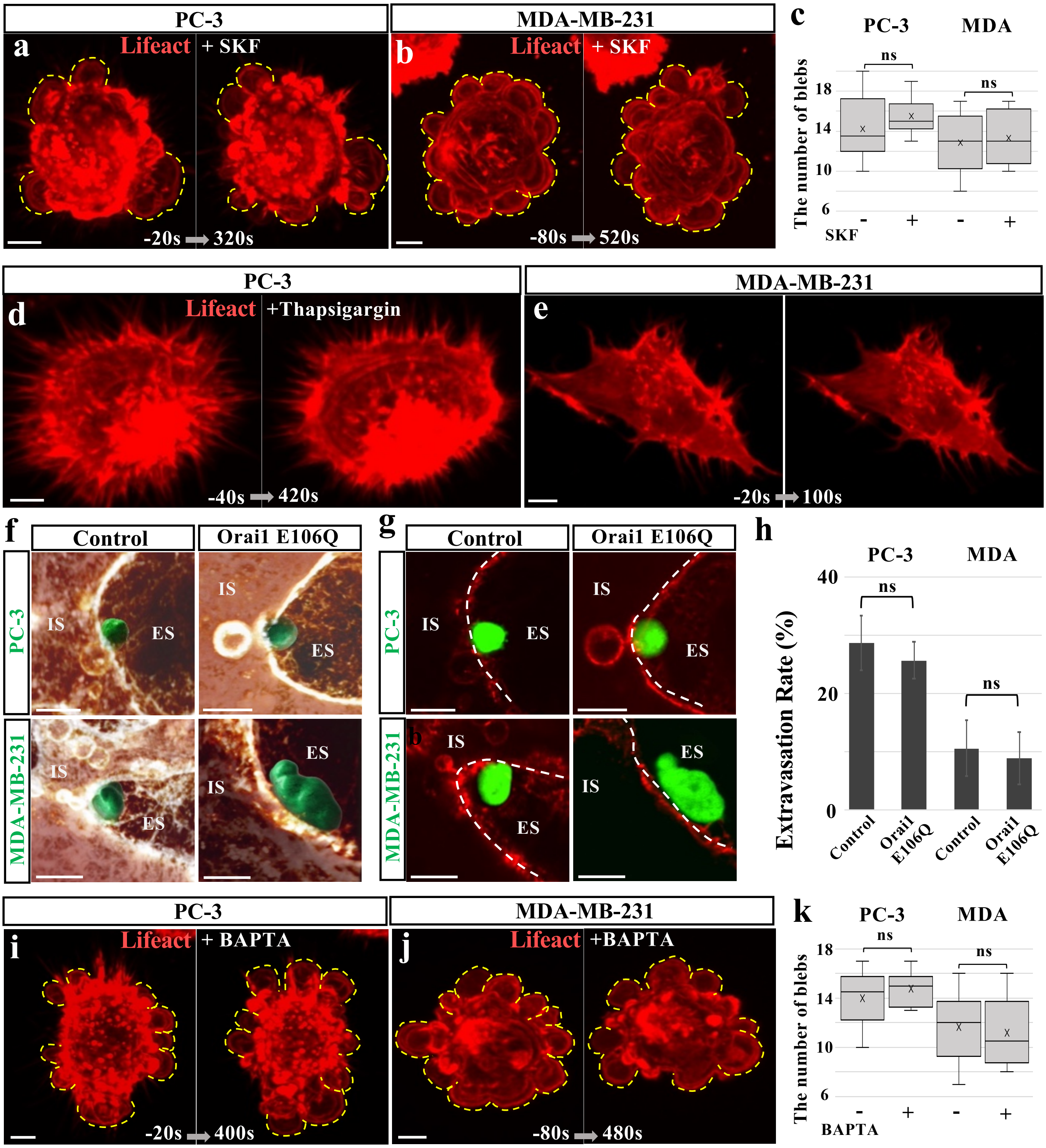


Extended Data Figure 9 Morita *et al.*

**Extended Data Figure 9.**
**Bleb formation and TEM of epithelial cancer cells are SOCE-independent.**
**a, b,** Time-lapse imaging of PC-3 (**a**) and MDA-MB-231 (**b**) cells exposed to SKF96365 in 2D culture. Time is shown in seconds relative to the initial addition of SKF96365. Yellow dotted lines outline the blebs. **c,** Quantification of bleb numbers before and after SKF96365 treatment in PC-3 and MDA-MB-231 ($N$=8 and 6 cells, respectively). **d, e,** Time-lapse imaging of PC-3 (**d**) and MDA-MB-231 (**e**) cells exposed to thapsigargin in 2D culture. **f,** 3D reconstructions of control and Orai1 E106Q-expressing PC-3 or MDA-MB-231 cells (green iso-surfaces) undergoing TEM in the YSV at 3.5 hpt. Blood vessels were labeled by intravenous injection of Rho-LCA (brown). Endothelial volumes were digitally clipped to visualize transmigrating cells. **g,** Images of transmigrated control or Orai1 E106Q-expressing PC-3 or MDA-MB-231 cell (green) in the YSV at 3.5 hpt. Blood vessels were visualized by injection of Rho-LCA (red). White dotted lines delineate intravascular (IS) and extravascular (ES) compartments. **h,** Quantification of extravasation efficiency for control and Orai1 E106Q-expressing PC-3 and MDA-MB-231 cells. Efficiency was calculated as the percentage of EGFP+ or Orai1 E106Q-expressing cells located outside the vasculature among all labeled cells in the YSV ($N$ = 4 embryos per group). **i, j,** Time-lapse imaging of PC-3 (**i**) and MDA-MB-231 (**j**) cells exposed to BAPTA in 2D culture. **k,** Quantification of the number of blebs before and after BAPTA treatment in PC-3 and MDA-MB-231 ($N$=8 and 6 cells, respectively). Error bars, s.e.m. ns, not significant (**c, k**: two-sided paired Student's $t$-test; **h**: Two-sided Welch's t-test). Scale bars: 5 μm in **a**, **b, d, e, i, j, n,** 20 μm in **f, g**, 2 μm in **i, j.**

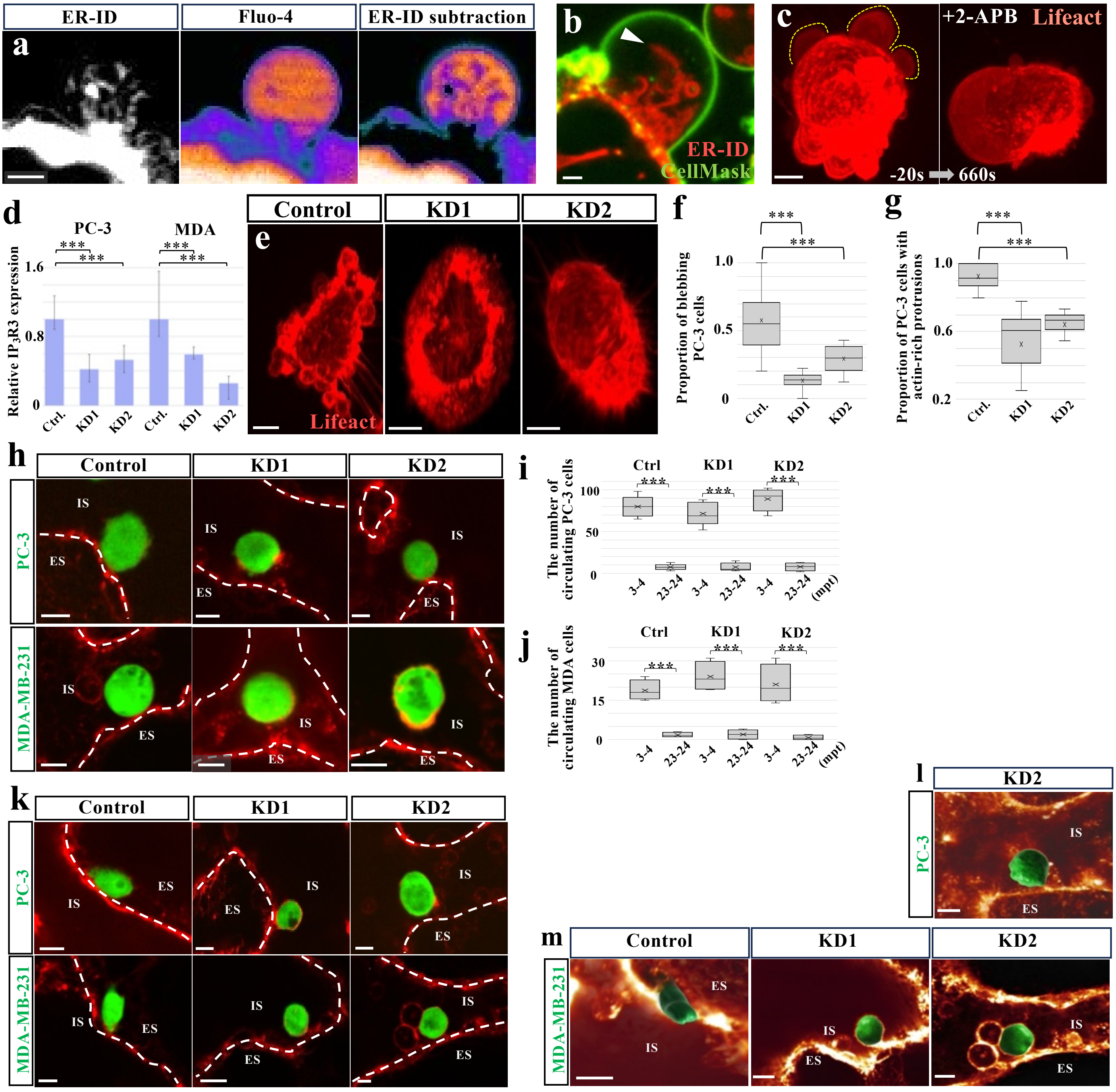


Extended Data Figure 10 Morita *et al.*

**Extended Data Figure 10.**

**$IP_3R$ is necessary for bleb formation and TEM of epithelial cancer cells.**

**a**, Dual imaging of ER-ID and Fluo-4 in a PC-3 cell during bleb formation. Left and middle: raw images; right: subtraction of ER–ID signal from Fluo-4. **b,** Dual visualization of CellMask (green) and ER-ID (red) showing ER tubules extending into a bleb of a PC-3 cell (white arrowhead). **c,** Time-lapse imaging of an MDA-MB-231 cell before and after 2-APB addition in 2D culture. Time is shown in seconds relative to the initial addition of 2-APB. The yellow dotted lines outline the emerging blebs. **d,** Knockdown efficiency of three different DsiRNAs targeting $IP_3R3$ in PC-3 and MDA-MB-231 cells. Relative expression levels were normalized to $IP_3R3$ expression in the negative control (set to 1). β-actin was used as a housekeeping gene. **e**, Representative images of Lifeact-expressing PC-3 cells: control, $IP_3R3$ KD1, and KD2, in 2D culture. **f,** Quantification of the proportion of bleb-forming PC-3 cells in 2D culture. **g,** Quantification of the proportion of PC-3 cells with actin-rich protrusions in 2D culture. **h,** Representative images of control, $IP_3R3$ KD1, and KD2 PC-3 and MDA-MB-231 cells in the YSV at 20 mpt. Blood vessels were visualized by injection of Rho-LCA (red). White dotted lines delineate the boundary between the intravascular space (IS) and the extravascular space (ES). **i, j,** Quantification of control, $IP_3R3$ KD1, and KD2 cells in PC-3 (**i;** *N*=6, 6, 4 embryos, respectively) and MDA-MB-231 (**j;** *N*=4 embryos, each) during passage through the YSV imaging field at 3–4 and 23-24 mpt. **k**, Representative images of control, $IP_3R3$ KD1, and KD2 PC-3 and MDA-MB-231 cells in the YSV at 3.5 hpt. **l, m**, 3D reconstructions of $IP_3R3$ KD2 PC-3 cells (**l**) and control, $IP_3R3$ KD1, and KD2 MDA-MB-231 cells (**m**) (green iso-surfaces) undergoing TEM in the YSV at 3.5 hpt. Blood vessels were labeled by intravenous injection of Rho-LCA (brown). Endothelial volumes were digitally clipped to visualize transmigrating cells. ns, not significant. *P<0.05, **P<0.01, ***P<0.001 (**d, f, g**: Two-sided Welch's t-test; **i, j**: two-sided, unpaired Student's *t*-test). Error bars represent standard deviation. Scale bars: 2 μm in **a-b,** 5 μm in **c, e** 10 μm in **h**, **k, l, m.**
