## Supplementary material for "Bleb-based extravasation: conserved morphodynamics, divergent calcium control": Table 1

**Expression of STIM/Orai related genes during male PGC development.**

| | Expression of genes (TPM*) in each embryonic stage | | |
|---|---|---|---|
| Gene/Transcipts ID# | E2.5 | E4.5 | E6.5 |
| Orai1/NM_001030658.2 | 50.35665 | 34.27645 | 31.8175 |
| Orai2/NM_001030710.3 | 34.99286 | 12.32332 | 7.32639 |
| STIM1/NM_001030838.3 | 0.2942275 | 1.3044365 | 0.4004635 |
| STIM2/XM_420749.8 | 11.82875 | 2.871365 | 3.206615 |

### NCBI transcripts ID based on genomic assembly (GCF_016699485.2). The major isoforms of each gene were selected.

* TPM was cullculated based on published datasets (GSE188689)

**Expression of STIM/Orai related genes during female PGC development.**

| | Expression of genes (TPM*) in each embryonic stage | | |
|---|---|---|---|
| Gene/Transcipts ID# | E2.5 | E4.5 | E6.5 |
| Orai1/NM_001030658.2 | 51.4785 | 32.74455 | 29.53685 |
| Orai2/NM_001030710.3 | 6.58821 | 6.049065 | 9.29929 |
| STIM1/NM_001030838.3 | 0.2192575 | 0.7551785 | 0 |
| STIM2/XM_420749.8 | 5.467405 | 4.631605 | 2.54444 |

### NCBI transcripts ID based on genomic assembly (GCF_016699485.2). The major isoforms of each gene were selected.

* TPM was cullculated based on published datasets (GSE188689)

**Informatics analysis.**

The published datasets (GSE188689) was used to investigate expression of STIM/Orai related genes.

Quality check and trimming of sequenced reads was performed using Trim Galore (ver.0.6.10) with FastQC (ver.0.11.9) and Cutadapt (ver.3.2).

Treated reads were mapped to transcripts (GCF_016699485.2) and TPM were caluculated using Kallisto (ver.0.46.0).
